# The meningeal transcriptional response to traumatic brain injury and aging

**DOI:** 10.1101/2022.06.16.496485

**Authors:** Ashley C. Bolte, Daniel A. Shapiro, Arun B. Dutta, Wei Feng Ma, Katherine R. Bruch, Ana Royo Marco, John R. Lukens

**Author notes:** These authors contributed equally to the manuscript (Ashley C. Bolte, Daniel A. Shapiro). **Correspondence should be addressed to:** John R. Lukens, Department of Neuroscience, Center for Brain Immunology and Glia, University of Virginia, 409 Lane Road, MR4-6154, Charlottesville VA 22908, Ashley C. Bolte, Department of Neuroscience, Center for Brain Immunology and Glia, University of Virginia, 409 Lane Road, MR4-6102, Charlottesville VA 22908.

## Abstract

Emerging evidence suggests that the meningeal compartment plays instrumental roles in various neurological disorders and can modulate neurodevelopment and behavior. While this has sparked great interest in the meninges, we still lack fundamental knowledge about meningeal biology. Here, we utilized high-throughput RNA sequencing (RNA-seq) techniques to investigate the transcriptional response of the meninges to traumatic brain injury (TBI) and aging in the sub-acute and chronic time frames. Using single-cell RNA sequencing (scRNA-seq), we first explored how mild TBI affects the cellular and transcriptional landscape in the meninges in young mice at one week post-injury. Then, using bulk RNA sequencing, we assessed the differential long-term outcomes between young and aged mice following a TBI. In our scRNA-seq studies, we found that mild head trauma leads to an activation of type I interferon (IFN) signature genes in meningeal macrophages as well as the mobilization of multiple distinct sub-populations of meningeal macrophages expressing hallmarks of either classically activated or wound healing macrophages. We also revealed that dural fibroblasts in the meningeal compartment are highly responsive to TBI, and pathway analysis identified differential expression of genes linked to various neurodegenerative diseases. For reasons that remain poorly understood, the elderly are especially vulnerable to head trauma, where even mild injuries can lead to rapid cognitive decline and devastating neuropathology. To better understand the differential outcomes between the young and the elderly following brain injury, we performed bulk RNA-seq on young and aged meninges from mice that had received a mild TBI or Sham treatment 1.5 months prior. Notably, we found that aging alone induced massive upregulation of meningeal genes involved in antibody production by B cells and type I IFN signaling. Following injury, the meningeal transcriptome had largely returned to its pre-injury signature in young mice. In stark contrast, aged TBI mice still exhibited massive upregulation of immune-related genes and markedly reduced expression of genes involved in extracellular matrix remodeling and maintenance of cellular junctions. Overall, these findings illustrate the dynamic and complex transcriptional response of the meninges to mild head trauma. Moreover, we also reveal how aging modulates the meningeal response to TBI.

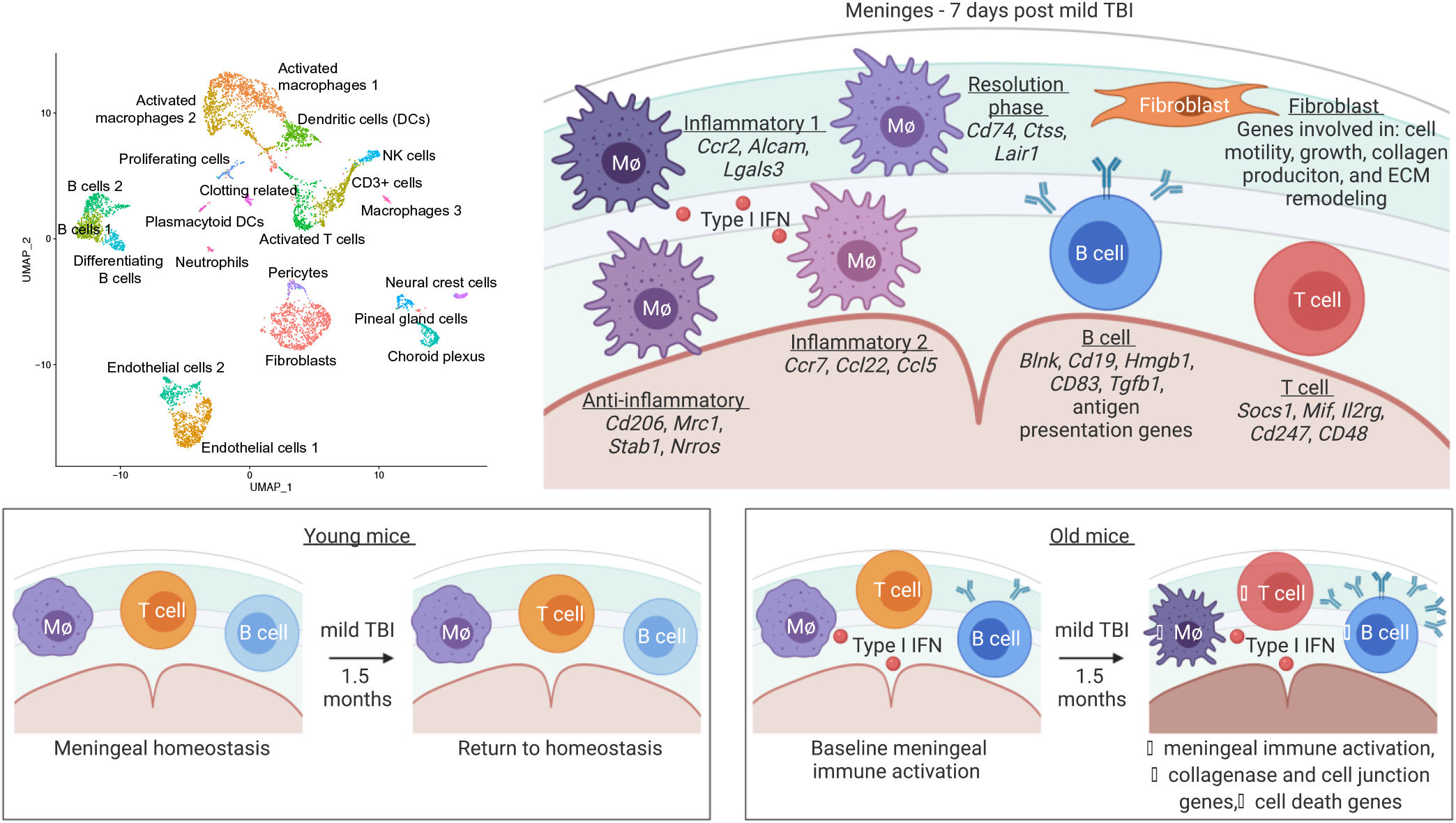

## BACKGROUND

Traumatic brain injury (TBI) affects millions of people each year and can result in devastating long-term outcomes (1–10). While TBI affects individuals of all ages, the elderly experience more severe consequences than do younger individuals with similar injury severity (11). The reason for this differential response to brain injury with respect to aging is not fully understood. Multiple findings have indicated that prolonged activation of the immune system following TBI may contribute to some of the negative TBI-associated sequelae (12–19). Interestingly, several studies point to differences in the immune response in elderly individuals that may contribute to more severe consequences following injury (17, 20–26). However, our understanding of the disparate central nervous system (CNS) responses between elderly and young individuals following TBI is still in its infancy.

Recent findings have implicated the meninges, a tri-layered tissue that resides between the brain parenchyma and skull, as an early responder to TBI and as a pivotal contributor to the CNS immune response following injury (27, 28). Meningeal enhancement with post-contrast fluid attenuated inversion magnetic resonance imaging (MRI) can be seen in 50% of patients with mild TBIs and no apparent parenchymal damage (28). This enhancement has been shown to occur within minutes of injury (29). Moreover, many individuals who experienced mild TBIs still exhibited extravasation of contrast into the sub-arachnoid space, indicating that the blood-brain-barrier was compromised (29). While most patients experienced resolution in meningeal enhancement 19 days after injury, about 15% had persistent enhancement three months post-injury, indicating that some patients experienced prolonged periods without complete meningeal repair following mild TBI (27). These protracted periods of meningeal enhancement likely represent ongoing inflammation within the compartment, yet the different cellular and molecular components that drive this inflammation have not been fully investigated.

The meningeal response to brain injury can be divided into several phases: acute, sub-acute, and chronic (28). Initial studies of the acute phase response after a mild TBI detail a meningeal response that consists of rapid meningeal cell death due to vascular leakage and reactive oxygen species release, which results in secondary parenchymal damage within the first several hours of injury (27, 28). The initial injury is followed by meningeal neutrophil swarming (present within an hour of injury) that is essential for regeneration of the initially damaged glial limitans (28). Disrupted meningeal vasculature is then repaired during the week following injury by non-classical monocytes (27). While these acute meningeal responses have been investigated, much less is understood about how brain injury shapes the meningeal environment more chronically, and if this response is affected by aging. Furthermore, it is unknown whether chronic meningeal changes following brain injury in aged individuals can contribute to neurodegenerative processes.

In addition to housing lymphatic vessels that drain molecules and cells to peripheral lymph nodes (30, 31), the meninges also contain a full array of innate and adaptive immune cells that are in constant communication with neurons and glia (32–34). In homeostasis, cytokine signaling from meningeal immune cells has been shown to be critical for shaping behavior (33, 35–37). For instance, IFN-γ is important in maintaining social behavior networks, whereas IL-4 production by meningeal T cells has been shown to influence learning and memory (35, 36). Recent studies also suggest that IL-17a secretion by γδ T cells in the meninges can impact anxiety-like behaviors and memory (33, 37). Of particular relevance, recent work has shown that an age-related decline of CCR7 expression by meningeal T cells may contribute to cognitive impairment, brain inflammation and neurodegenerative disease (38). In addition to meningeal T cell production of cytokines, it is known that immune cells within the cerebrospinal fluid (CSF) in the subarachnoid space can also produce signaling molecules and interact with brain-derived products and antigens (39, 40). Brain interstitial fluid (ISF) and CSF intermix in the subarachnoid space and both recirculate throughout the brain via the glymphatic system and drain through the meningeal lymphatic network to the periphery (30, 31, 41–50). This system provides meningeal cells and cells within the CSF with access to brain antigens and proteins. Despite mounting evidence demonstrating that meningeal cells can impact various aspects of neurobiology, we still lack a complete picture of how the meninges respond to processes that have been broadly linked to neurological disease, such as brain injury and aging. Likewise, little is known in regards to how aging impacts meningeal biology both under steady-state conditions and in response to TBI.

Here, we investigated how the meningeal transcriptional environment is altered following TBI and in aging utilizing high-throughput sequencing techniques, namely single-cell RNA sequencing (scRNA-seq) and bulk RNA sequencing (RNA-seq). We focused on sub-acute (one week post-TBI) and chronic (1.5 months post-TBI) time points after brain injury to better understand how the cellular makeup and gene expression profiles in the meninges change with time and with age. Furthermore, we aimed to reveal how chronic changes in the meningeal transcriptional landscape may contribute to, or be a product of long-term neurodegenerative changes. Herein, we find that the heterogeneous cellular makeup of the meninges is altered one week following TBI with an increase in the frequency of macrophages and fibroblasts. Moreover, we further show that the meningeal transcriptional environment is massively altered in aging, including a sweeping upregulation of genes involved in antibody production and type I interferon (IFN) signaling. When examining the genes that were upregulated in aged mice as compared to young mice 1.5 months following TBI, we found that there is a broad downregulation in genes important for extracellular matrix remodeling and collagen production, and an overall activation of the immune system. This prolonged activation of the immune system is unique to the aged TBI mice, as young mice exhibit few alterations in the meningeal transcriptome 1.5 months following injury. These findings highlight the dynamic nature of the meningeal transcriptome in response to TBI and aging, and shed light on some of the differences between young and aged individuals in responding to brain injury.

## METHODS

### Mice

All mouse experiments were performed in accordance with the relevant guidelines and regulations of the University of Virginia and were approved by the University of Virginia Animal Care and Use Committee. Young C57BL/6J wild-type (WT) mice were obtained from Jackson Laboratories. All WT aged mice were approximately 20 months of age and were obtained from the National Institute on Aging (NIA) Aged Rodent Colonies. The mice from the NIA Aged Rodent Colonies were originally derived from the Jackson colonies. Upon arrival, aged mice were housed in University of Virginia facilities for at least 2 months before use. Other young mice ordered directly from the Jackson Laboratories were housed in the local University of Virginia facility for at least 2 weeks before use. Mice were housed in specific pathogen-free conditions under standard 12-hour light/dark cycle conditions in rooms equipped with control for temperature (21 ± 1.5°C) and humidity (50 ± 10%). Male mice were used for all studies.

### Traumatic brain injury

This injury paradigm was adapted from the published Hit and Run model (51, 52). Mice were anesthetized by 4% isoflurane with 0.3 kPa O_2_ for 2 minutes and then the right preauricular area was shaved. The mouse was placed prone on an 8 x 4 x 4-inch foam bed (type E bedding, open-cell flexible polyurethane foam with a density of approximately 0.86 pounds per cubic feet and a spring constant of approximately 4.0 Newtons per meter) with its nose in a nosecone delivering 1.5% isoflurane (purchased from Foam to Size, Ashland VA). The head was otherwise unsecured. The device used to deliver TBI was a Controlled Cortical Impact Device (Leica Biosystems, 39463920). A 3 mm impact probe was attached to the impactor device which was secured to a stereotaxic frame and positioned at 45 degrees from vertical. In this study, we used a strike depth of 2 mm, 0.1 s of contact time and an impact velocity of 5.2 meters (m) per second (s). The impactor was positioned at the posterior corner of the eye, moved 3 mm towards the ear and adjusted to the specified depth using the stereotaxic frame. A cotton swab was used to apply water to the injury site and the tail in order to establish contact sensing. To induce TBI, the impactor was retracted and dispensed once correctly positioned. The impact was delivered to the right inferior temporal lobe of the brain. Following impact, the mouse was placed supine on a heating pad and allowed to regain consciousness. After anesthesia induction, the delivery of the injuries took less than 1 minute. The time until the mouse returned to the prone position was recorded as the righting time. Upon resuming the prone position, mice were returned to their home cages to recover on a heating pad for six hours with soft food. For Sham procedures, mice were anesthetized by 4% isoflurane with 0.3 kPa O_2_ for 2 minutes and then the right preauricular area was shaved. The mouse was placed prone on a foam bed with its nose secured in a nosecone delivering 1.5% isoflurane. The impactor was positioned at the posterior corner of the eye, moved 3 mm towards the ear and adjusted to the specified depth using the stereotaxic frame. A cotton swab was used to apply water to the injury site and the tail in order to establish contact sensing. Then, the impactor was adjusted to a height where no impact would occur, and was retracted and dispensed. Following the Sham procedure, the mouse was placed supine on a heating pad and allowed to regain consciousness. Mice were allowed to recover on the heating pad in their home cages for 6 hours with soft food before being returned to the housing facilities.

### Tissue collection

Mice were euthanized with CO_2_ and then transcardially perfused with 20 ml 1x PBS. For all meningeal collections, the meninges on the skullcap dorsal to the ear canal were collected. The dorsal meninges do not include the direct site impacted by the TBI. For meningeal whole mount preparation, the skin and muscle were stripped from the outer skull and the skullcap was removed with surgical scissors and fixed in 2% PFA for 12 hours at 4°C. Then the meninges (dura mater and some arachnoid mater) were carefully dissected from the skullcaps with Dumont #5 forceps (Fine Science Tools). Meningeal whole-mounts were then moved to PBS and 0.05% sodium azide in PBS at 4 °C until further use. For meninges collection for scRNA-seq, the skullcap was removed as previously described, and placed into DMEM medium. Meninges were then scraped from the skullcap and processed further to create a single cell suspension as described in the method’s section entitled ‘Meningeal preparation for scRNA-seq’ below. For meninges collection for bulk RNA-seq, the skullcap was removed as described above, and placed into DMEM medium. The meninges were scraped from the skullcap and were immediately snap-frozen at −80°C in TRIzol (15596018, Life Technologies), until further use. Brains were removed and kept in 4% PFA for 24 h and then cryoprotected with 30% sucrose for 3 days. A 4 mm coronal section of brain tissue that surrounded the site of injury was removed using a brain sectioning device and then frozen in Tissue-Plus OCT compound (Thermo Fisher Scientific). Fixed and frozen brains were sliced (50 -μm thick sections) with a cryostat (Leica) and kept in PBS + 0.05% sodium azide in PBS at 4 °C until further use.

### RNA extraction and sequencing

For RNA extraction from the meningeal tissue, the meninges were harvested as described in the ‘Tissue Collection’ methods section above and snap-frozen in 500μL TRIzol Reagent (15596018, Life Technologies) and stored at −80°C until further use. For each of the four experimental groups (Young Sham, Aged Sham, Young TBI and Aged TBI) 2 dorsal meningeal samples were combined to create 1 biological replicate. Three biological replicates were used for each experimental group yielding a total of 12 samples comprised of 2 meninges each. After defrosting on ice, 10 silica beads were added to each tube and the tissue was homogenized for 30 seconds using a mini bead beater. Following homogenization, the samples were centrifuged for 12,000 xg for 10 minutes at 4°C. The supernatant was transferred to a new tube and incubated at room temperature for 5 minutes. Next, 0.1 mL of chloroform was added to the supernatant, vortexed, incubated for 2 minutes at room temperature and then centrifuged at 12,000 xg for 15 minutes at 4°C. The top aqueous phase was transferred into a new Eppendorf tube and the RNeasy Micro Kit (74004, Qiagen) was used to isolate the RNA. RNA was frozen at −80°C until sent for sequencing. For sequencing, total RNA samples were sent to GENEWIZ for library preparation and paired-end sequencing.

### Meningeal preparation for scRNA-seq

The day before meningeal harvest, Eppendorf tubes were coated with FACS buffer (1% BSA, 1mM EDTA in PBS) overnight. Mice were euthanized with CO_2_ and then transcardially perfused with ice-cold PBS with heparin (0.025%). The skull caps were prepared as described in ‘Tissue Collection’. Meninges were peeled from the skull cap and placed in ice-cold DMEM for the entirety of collection. The meninges from 5 mice that had received TBI 1 week prior were pooled as one biological replicate. The meninges from 5 mice that had received a Sham procedure 1 week prior were pooled as one biological replicate. These 2 samples were then processed for scRNA sequencing. Meninges were then digested for 15 minutes at 37°C with constant agitation using 1 mL of pre-warmed digestion buffer (DMEM, with 2% FBS, 1 mg/mL collagenase VIII (Sigma Aldrich), and 0.5 mg/mL DNase I (Sigma Aldrich)). The enzymes were neutralized with 1 mL of complete medium (DMEM with 10% FBS) and meninges were then filtered through a 70 µm cell strainer. An additional 2 mL of FACS buffer was added, samples were centrifuged at 400 xg for five minutes, and samples were resuspended in FACS buffer. After 2 washes, cells were resuspended in FACS buffer with DAPI (0.2 µg/mL). Singlet gates were selected using pulse width of the side scatter and forward scatter. Live cells were selected based on the lack of DAPI staining. Cells were sorted into 1.5 mL tubes with ice cold DMEM. Following sorting, cells were centrifuged again at 450 xg for 4 mins and the media was aspirated. Cells were resuspended in 200 μL 0.04% BSA in PBS ( 0.04% non-acetylated BSA) and centrifuged again. Cells were counted in 20 μL of 0.04% BSA in PBS using trypan blue. Approximately 4,000 cells per sample were loaded onto a 10X Genomics Chromium platform to generate cDNAs carrying cell- and transcript-specific barcodes and sequencing libraries constructed using the Chromium Single Cell 3’ Library & Gel Bead Kit 2. Libraries were sequenced on the Illumina NextSeq using pair-ended sequencing, resulting in 50,000 reads per cell.

### scRNA-seq analysis

The raw sequencing reads (FASTQ files) were aligned to the Genome Research Consortium (GRC) mm10 mouse genome build using Cell Ranger (v1.3.1) which performs alignment, filtering, barcode counting and unique molecular identifier (UMI) counting. RStudio (v1.2.5033) was used for all downstream analyses and Seurat (v.3.9.9) was used for filtering out low-quality cells, normalization of the data, determination of cluster defining markers and graphing of the data on UMAP (53, 54). Only one sequencing run was performed therefore there was no need for batch correction. Initially, there were 2261 cells collected from the Sham mice and 4022 cells collected from the TBI mice. Low-quality cells were excluded in an initial quality-control (QC) step by removing cells with fewer than 150 unique genes and cells expressing more than 5,000 unique genes in effort to remove doublets and triplets (Sham total: 2257, TBI total: 4018). Cells with transcriptomes that were more than 20% mitochondrial-derived were removed and cells with more than 5% hemoglobin among their expressed genes were also removed (Sham total: 2049, TBI total: 3775). Using Seurat, genes with high variance were selected using the FindVariableGenes() function, then the dimensionality of the data was reduced by principle component analysis (PCA) and identified by random sampling of 20 significant principal components (PCs) for each sample with the PCElbowPlot() function. Cells were clustered with Seurat’s FindClusters() function. Absolute cell counts for each population can be found in Table 1. scCATCH (v2.1) was used for automated cluster naming (55), and all cluster names were manually checked due to the lack of literature regarding meningeal cell populations. Next, differential gene expression analysis was performed within the clusters using the ZINB-WaVE (v1.12.0) and DESeq2 (v1.30.0) packages (56). Cytoscape (v3.8.0) and ToppCluster (https://toppcluster.cchmc.org/) were used for network analyses (57, 58). Data was organized and graphs were created using ggplot2, tidyverse, treemapify, circlize, Seurat and dplyr (59, 60). Pseudotime analysis was conducted using Monocle3 (v0.2.3.0) (61). All code used for analysis is available upon request.

**Table 1.** Counts of each cell population separated by Sham and TBI. Male WT mice at 10 weeks of age received a TBI or Sham procedure. One week later, the meninges from 5 mice per group were harvested, pooled, and processed for scRNA-seq. The cell counts for each cell population are shown after data processing.

### Bulk RNA-seq analysis

The raw sequencing reads (FASTQ files) were aligned to the GRC mm10 mouse genome build using the splice-aware read aligner HISAT2 (62). Samtools was used for quality control filtering (63). Reads were sorted into feature counts with HTSeq (64). DESeq2 (v1.30.0) was used to normalize the raw counts based on read depth, perform principal component analysis, and conduct differential expression analysis= (65). The p-values were corrected with the Benjamini-Hochberg procedure to limit false positives arising from multiple testing. The gene set collections from MSigDB were used for differential gene set enrichment analysis (66). The analysis itself was performed using the Seq2Pathway, fgsea, tidyverse, and dplyr software packages. Heatmaps were generated using the pheatmap R package while other plots were made with the lattice or ggplot2 packages. All code used for analysis is available upon request.

### Immunohistochemistry, imaging, and quantification

For immunofluorescence staining, meningeal whole mounts and floating brain sections in PBS and 0.05% sodium azide were blocked with a solution containing 2% goat serum or 2% donkey serum, 1% bovine serum albumin, 0.1% triton, 0.05% tween-20, and 0.05% sodium azide in PBS for 1.5 h at room temperature (RT). This blocking step was followed by incubation with appropriate dilutions of primary antibodies in blocking solution at 4°C overnight. The primary antibodies and their dilutions include: anti-Collagen I (Abcam, ab21286, 1:200), anti-J chain (Invitrogen, SP105, 1:200), anti-Lyve-1-efLuor 488 (eBioscience, clone ALY7, 1:200), anti-Iba1 (Abcam, ab5076, 1:300), anti-GFAP (Thermo Fisher Scientific, 2.2B10, 1:1000), anti-MHC Class II 660 (eBioscience, M5/114.15.2, 1:100), anti-Ifnar1 (Thermo Fisher Scientific, SR45-08, 1:150), anti-CD31 (Millipore Sigma, MAB1398Z, clone 2H8, 1:200), anti-B220 (Thermo Fisher Scientific, RA3-6B2, 1:200) and anti-NeuN (EMD Millipore, Mab277, clone A60, 1:500). Meningeal whole mounts and brain sections were then washed three times for 10 min at room temperature in PBS and 0.05% tween-20, followed by incubation with the appropriate goat or donkey Alexa Fluor 488, 594 or 647 anti-rat, -goat or -rabbit (Thermo Fisher Scientific, 1:1000) or - Armenian hamster (Jackson ImmunoResearch, 1:1000) IgG antibodies for 2h at RT in blocking solution. The whole mounts and brain sections were then washed 3 times for 10 min at RT before incubation for 10 min with 1:1000 DAPI in PBS. The tissue was then transferred to PBS and mounted with ProLong Gold antifade reagent (Invitrogen, P36930) on glass slides with coverslips.

Slide preparations were stored at 4 °C and imaged using a Lecia TCS SP8 confocal microscope and LAS AF software (Leica Microsystems) within one week of staining. Quantitative analysis of the acquired images was performed using Fiji software and Imaris software. Imaging parameters for brightness, contrast, and threshold values were applied uniformly throughout each experiment. Additionally, tears in the meninges were excluded when performing the analyses. For assessment of IFNAR1+Iba1+ number and volume in meningeal whole mounts, 5 meningeal whole mounts were included per experimental group. Images at 63x were taken at four uniform positions along the transverse sinus (2 on each side of the confluence of the sinuses). Number and volume of IFNAR1+ puncta colocalized with Iba1+ cells was calculated for each image using Imaris. All four images were averaged together to calculate the average number or volume of puncta per mouse. For the assessment of collagen expression and J-chain quantification, 5 meningeal whole mounts were included per experimental group. For collagen, 10x images were taken of meningeal whole mounts. To quantify the collagen, the corrected total cellular fluorescence (CTCF) was used, which takes into account the area of the meninges, the average fluorescent intensity of that area, and the fluorescent intensity of the background, as given by the formula: Mean fluorescence of meningeal whole mounts - (area of meningeal whole mount x mean fluorescence of background). For J-chain quantification, a uniformly sized 20x image of the superior sagittal sinus was taken for each sample with 5 samples in each experimental group. The number of J-chain puncta was then quantified (puncta threshold: 5-10 microns) using the “Analyze Particle” tool. For quantification of B220+ cells, 20x scans (n=5 per group) were taken of the entire transverse sinus. The number of B220+ cells were manually counted and quantified by a blinded experimenter. For CD31 staining, percent area was quantified from whole mount meninges scanned at 10x. For MHC II staining, eight meningeal whole mounts per experimental group were imaged at 10x. The number of MHC II puncta was quantified after a threshold of 200-400 microns squared had been applied.

For the assessment of gliosis in the injured and uninjured brains in Supplementary Fig. 1, two representative brain sections from the site of the lesion (approximately −0.74 to 0 bregma) or the corresponding area in Sham animals were fully imaged and at least 4 animals were included per experimental group. The full brain section was adjusted for brightness and contrast uniformly for each experiment and each hemisphere was traced, and then the threshold was uniformly set for each experiment to select for stained cells. The percent area of coverage of each immunohistochemical markers was calculated for the hemisphere ipsilateral to the injury site (right) for each brain section. The mean percent area fraction was calculated using Microsoft Excel. For Supplementary Fig. 1, high magnification images (63x) were taken directly adjacent to the site of the injury.

### Statistical analysis and reproducibility

Sample sizes were chosen on the basis of standard power calculations (with *α* = 0.05 and power of 0.8). Experimenters were blinded to the identity of experimental groups from the time of euthanasia until the end of data collection and analysis. Statistical analysis was performed using RStudio (v1.2.5033) and GraphPad Prism (v8.4.3). Individual statistical analyses for each experiment are indicated in the corresponding figure legends.

### Data availability

All data and genetic material used for this paper are available from the authors on request.

### Code availability

All code used for analysis is available at [https://github.com/danielshapiro1/MeningealTransciptome] or upon request.

## RESULTS

### Mild TBI incites alterations in the cellular composition of the meninges

To gain insights into how TBI impacts meningeal biology, we subjected mice to a mild closed-skull injury and then performed scRNA-seq on the meninges one week post-injury (Figure 1a). In this model of mild TBI, mice received a single hit to the right inferior temporal and frontal lobes using a stereotaxic electromagnetic impactor (Supplementary Figure 1a) (52). Of note, we have previously shown that head injury in this model does not result in appreciable alterations in balance, motor coordination, reflex, and alertness (52). Consistent with the mild nature of this TBI model, we also do not observe any appreciable differences in CD31 blood vasculature staining at 24 hrs following head trauma (Supplementary Figure 1b,c). Moreover, we only detect modest increases in gliosis (Iba1 and GFAP staining) (Supplementary Figure 1d,e,f) and MHCII+ staining in the meninges at 24 hrs post-TBI (Supplementary Figure 1g,h,i).

**Figure 1.**
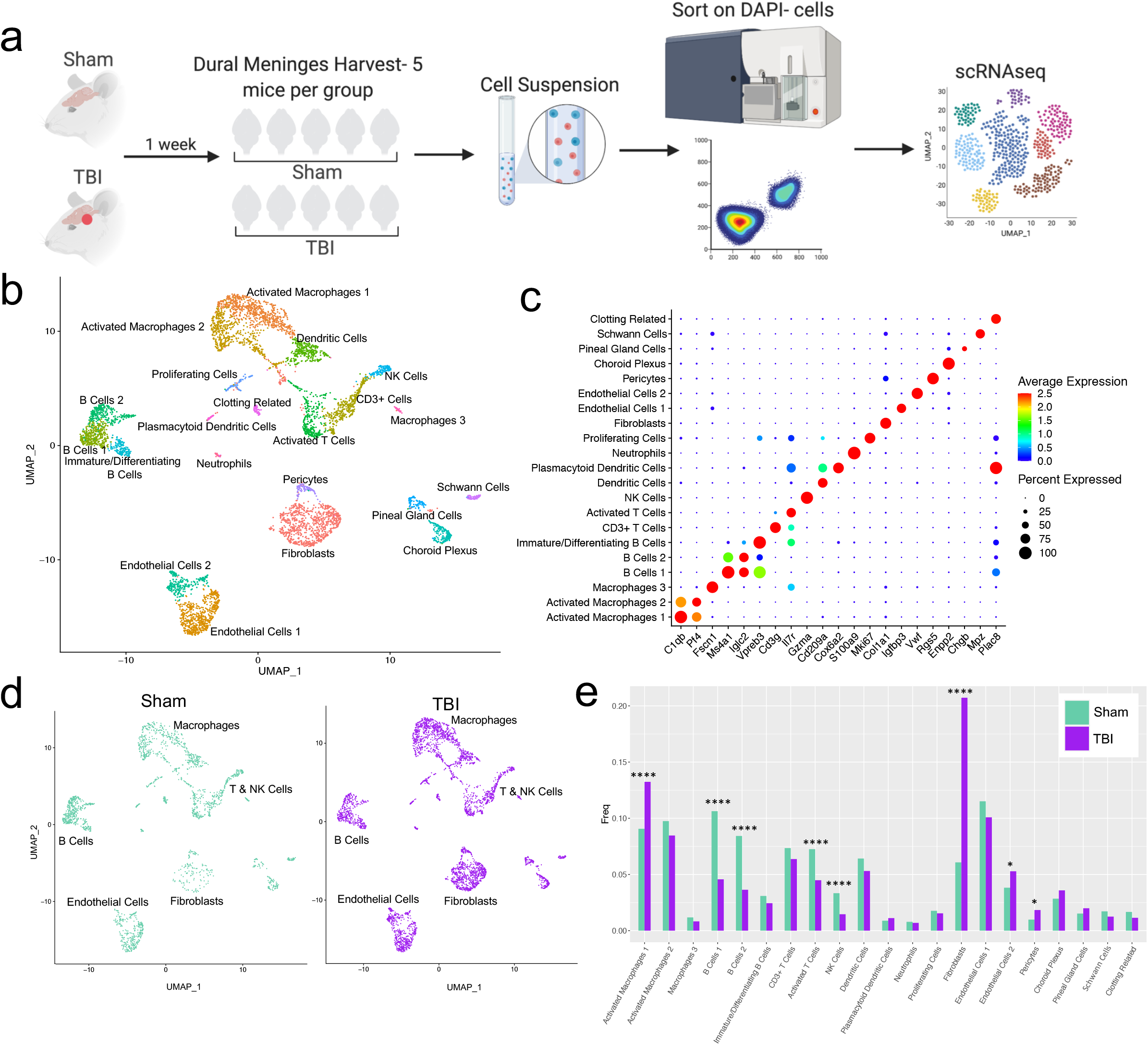
Alterations in the composition of meningeal cell populations following brain injury. Male C57BL/6J wild-type (WT) mice at 10 weeks of age were subjected to a mild closed-skull injury above the right inferior temporal lobe or Sham procedure. One week later, the meninges from 5 mice per group were harvested, pooled, and processed for scRNA-seq. a) Schematic of scRNA-seq protocol. b) Uniform Manifold Approximation and Projection (UMAP) representation of the cell populations present in the meninges where both Sham and TBI groups are included. Colors are randomly assigned to each cell population. c) Dot plot representation of cluster defining genes for each cell population, where each gene represents the most significant cluster-defining marker for each population. The color and size of each dot represents the average expression and percent of cells expressing each gene, respectively. d) UMAP representations of the cell populations present in the meninges separated by Sham (sage) and TBI (purple). e) Frequencies of cell populations in Sham vs. TBI samples represented as a gradient bar chart. Graphs were calculated using Seurat by normalizing the dataset, finding the variable features of the dataset, scaling the data, and reducing the dimensionality. Each data point in a UMAP plot represents a cell. P values were calculated using a two sample z-test. ****P<0.0001, *P<.05, bar chart pairs without * were not statistically significant.

For all of our sequencing studies in this paper, we strategically chose to isolate only the dorsal meningeal tissue, as this region of the meninges does not include the tissue affected by the direct injury site. Therefore, the sequencing data generated from these studies should better reflect the global meningeal changes that result from a localized injury site rather than the tissue damage and response at the immediate injury site. Joint clustering of both the Sham and TBI meninges revealed 21 unique cell populations including endothelial cells, fibroblasts, Schwann cells, and ciliated ependymal cells from the pia (Figure 1b,c, Table 1, Supplementary Figure 2). Additionally, the meninges contained a full repertoire of immune cells including macrophages, B cells, T cells, NK cells, dendritic cells, plasmacytoid dendritic cells, and neutrophils (Figure 1b,c, Table 1, Supplementary Figure 2). Other cell populations were less well-defined and included cells expressing genes important for clotting and proliferating cells (Figure 1b,c, Table 1, Supplementary Figure 2). When separated out by Sham and TBI treatments, all 21 populations were still present in both groups (Figure 1d, Table 1), however the frequencies were varied (Figure 1e). Following brain injury, there was a higher frequency of one group of macrophages which we denoted as “Activated Macrophages 1” as they exhibit high expression of complement-related genes (Figure 1c,e). Moreover, the frequency of fibroblasts was substantially increased following head trauma (Figure 1e). While there was a reduction in frequency of some other cell types, namely the B cell populations, it is unclear whether this was relative to the expansion of the other subsets or an actual decrease in number (Figure 1e). In order to ensure the short digestion and processing steps of the sample preparation did not result in significant upregulation of stress-related genes in both Sham and TBI samples, we examined a collection of genes that have been known to be upregulated after tissue processing and in stress-related conditions (67–69) (Supplementary Figure 3). Very few sequenced cells expressed these genes and there were not substantial differences between the TBI or Sham experimental groups, suggesting minor contributions of processing on gene expression and similar effects across experimental groups (Supplementary Figure 3). Overall, these data highlight the heterogeneous nature of the meningeal tissue and also demonstrate that the frequencies of macrophage and fibroblast populations are increased one week post-TBI.

### Effects of mild TBI on the meningeal macrophage transcriptome

Given our data demonstrating an appreciable expansion of the meningeal macrophage population following injury (Figure 1e) as well as emerging data suggesting instrumental roles for these cells in TBI pathogenesis (27, 28), we decided to focus first on the response of meningeal macrophages to head trauma (Figure 1e). Differential gene expression analysis of the “Activated Macrophage” populations (Activated Macrophages 1 & 2) demonstrated 321 upregulated genes and 369 downregulated genes following head injury when using a false discovery rate of <0.1 (Figure 2a). When we performed a network analysis on the significantly upregulated genes from these populations, we found an enrichment of pathways related to immune system activation. Upregulated genes in the activated population included those important for cytokine secretion, immune cell differentiation, motility, and chemotaxis (Figure 2b). Furthermore, the most highly enriched gene ontology (GO) biological processes modulated in response to head trauma were found to be related to immune system activation and the stress response (Figure 2c). We also noticed that some of the most significantly upregulated genes contributing to the immune-related GO terms were important for the type I IFN response including *Ifnar1, Ifi203, Irf2bp2,* and *Irf5*, amongst others (Figure 2d). At the protein level, we confirmed that 1 week after injury or a Sham procedure, there was a substantial increase in IFNAR1 expression by Iba1+ macrophages in the TBI group when compared to the Sham group when examining both the volume (Figure 2f) and number (Figure 2g) of IFNAR1+ puncta along the transverse sinus in high magnification images of meningeal whole mounts (Figure 2e,g). Interestingly, recent studies suggest that elevated type I IFN signaling in the brain parenchyma is a driver of detrimental outcomes in TBI pathogenesis (70, 71). Taken together, these findings suggest that meningeal macrophages upregulate inflammation-related genes one week following brain injury and may contribute to the type I IFN signature that is seen following TBI.

**Figure 2.**
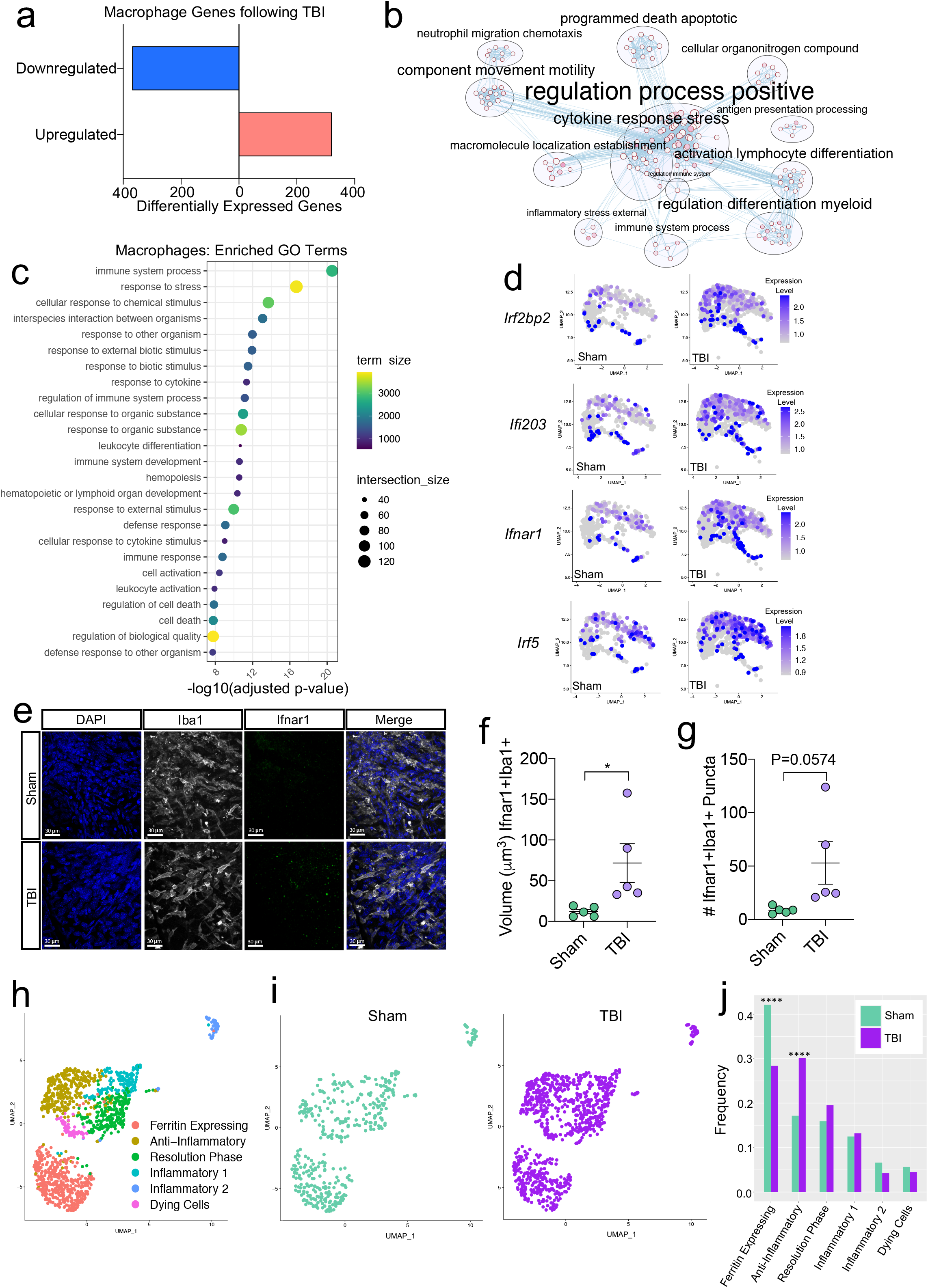
Transcriptional response of meningeal macrophages to mild TBI. Male WT mice at 10 weeks of age received a TBI or Sham procedure. One week later, the meninges from 5 mice per group were harvested, pooled, and processed for scRNA-seq. a) Quantification of the number of upregulated and downregulated macrophage genes following injury (FDR<0.1). b) Network analysis of significantly upregulated genes in meningeal macrophages following injury. Text size is proportional to the number of genes enriched in that cluster. Node size is roughly proportional to the number of GO terms in that cluster (node size was manually adjusted so may not be exactly proportional to GO terms included). Dot size is proportional to the number of genes contributing to each GO term. Dot color is proportional to p-value, where colors closer to white have lower p-values. Connecting lines represent GO terms with shared genes, more lines represents a higher number of shared genes between nodes. c) Dot plot showing the 25 most enriched GO terms with significantly upregulated genes following TBI in the meningeal macrophage population. The color and size of each dot represents the size of the GO term and the number of upregulated genes that contribute to each term, respectively. d) Feature plots depicting several significantly upregulated genes following injury (FDR<0.1). The color of each data point represents the expression level of the indicated gene within that cell. Mice at 10 weeks of age received a TBI or Sham procedure. One week later, the mice were harvested and meningeal whole mounts were processed for immunohistochemistry. e) Representative 63x images in Sham and TBI mice of cells along the transverse sinus stained for DAPI (blue), Iba1 (grey) and IFNAR1 (green). Quantification of the volume of IFNAR1+Iba1+ puncta (f) and number of IFNAR1+Iba1+ puncta in high magnification 63x images (g) along the transverse sinuses. Each data point represents an individual mouse. h) UMAP representation showing re-clustering of the meningeal macrophage populations. i) UMAP representation of the macrophages present in the meninges separated by Sham (sage) and TBI (purple). j) Frequencies of meningeal macrophage populations in Sham vs. TBI samples represented as a gradient bar chart. Graphs were calculated using Seurat by normalizing the dataset, finding the variable features of the dataset, scaling the data, and reducing the dimensionality. Differential gene expression was calculated using the ZINB-WaVE function for zero-enriched datasets and DESeq2. Each data point in a UMAP plot represents a cell. P values for (f-g) were calculated using unpaired two-sample students t-tests and P values for (j) were calculated using a two sample z-test. *P<0.05, ****P<0.0001. Bar chart pairings without * were not statistically significant. FDR; false discovery rate.

Previous findings also suggest that there are several subtypes of meningeal macrophages that respond to TBI (27). Therefore, we decided to look more closely at the subpopulations within the original macrophage clusters. We re-clustered the three macrophage populations (Activated Macrophages 1 & 2, and Macrophages 3) combined from Sham and TBI meninges, which yielded six different meningeal macrophage clusters (Figure 2h). The largest population of macrophages expressed high levels of ferritin (“Ferritin Expressing”) (Figure 2h, Supplementary Figure 4a). The top two cluster-defining genes within the “Ferritin Expressing” macrophages were ferritin light chain (Ftl1) and ferritin heavy chain (Fth1) (Figure 2h, Supplementary Figure 4a). There were two additional populations, deemed ‘Anti-Inflammatory’ and ‘Resolution Phase’ macrophages, that appeared to be alternatively activated, anti-inflammatory macrophages that are likely implicated in the healing response following injury. The top cluster-defining gene in the “Anti-Inflammatory” macrophage cluster was *Mrc1* (also referred to as CD206), which is known to be present on macrophages that play a role in the healing response after TBI (27). Other highly-significant subcluster-defining genes in the “Anti-Inflammatory” macrophage population included *Stab1, Nrros*, and *Dab2*, which are known to be expressed on healing macrophages, and are important for repressing reactive oxygen species and limiting type I IFN responses (72–74) (Supplementary Figure 4a,b). “Resolution Phase” macrophages do not fall into either the M1 classically activated or M2 alternatively activated macrophage categories and are believed to play a regulatory role following an inflammatory event (75). They tend to be enriched for antigen presenting genes, chemokine genes, and proliferation-related genes (75). Indeed, the meningeal macrophages in the “Resolution Phase” cluster were defined by their expression of antigen presentation-related genes (*H2-Eb1, H2-Ab1, H2-Aa, Cd74, Ctss*) and anti-inflammatory genes such as *Lair1*, an inhibitory receptor that prevents over-activation of cytokine production (76) (Supplementary Figure 4a,c). In contrast to these “Anti-Inflammatory” and “Resolution Phase” clusters, the final two macrophage populations exhibited gene signatures more commonly associated with inflammatory macrophages (Figure 2h). The “Inflammatory 1” macrophage cluster was defined by its differential expression of *Ccr2* and adhesion molecules such as *Alcam* and *Lgals3* (Supplementary Figure 4a,d). The second inflammatory macrophage subset “Inflammatory 2” was defined by its expression of genes important for chemotaxis including *Ccr7, Ccl22*, and *Ccl5* (77, 78) (Supplementary Figure 4a,e).

To determine how injury affected these distinct meningeal macrophage populations, we separated the cells into Sham and TBI groups and examined their frequencies (Figure 2i,j). Interestingly, there was an overall relative increase in the “Anti-Inflammatory” and “Resolution Phase” macrophages in the TBI group, indicating that one week after injury, the response of the meningeal macrophages appears to shift towards wound healing and inflammation resolution (Figure 2i,j). There was also a relative reduction in the “Ferritin Expressing” macrophages following injury (Figure 2i,j). Overall, these findings demonstrate that although the macrophages collectively upregulated genes essential for inflammation following TBI, the frequencies of “Resolution Phase” and “Anti-Inflammatory” macrophages also increased and may play a role in the wound healing process.

### Mild TBI induces the upregulation of neurological disease-associated genes in meningeal fibroblasts

Next, we wanted to further investigate the “Fibroblast” population, as it was expanded 1 week following injury in the single cell dataset (Figure 1d,e). After head injury, the “Fibroblast” population exhibited 368 activated genes and 320 repressed genes (Figure 3a). There were several patterns in the activated genes following injury, including genes important for collagen production and extracellular matrix remodeling (*Nisch, Ppib, Pmepa1, Ddr2*) and genes critical for cell motility and growth (*Ptprs, Pfn1, Cd302, Tpm3*) (Figure 3b). To validate these sequencing findings, we utilized immunohistochemical staining for collagen, which is produced by fibroblasts, in meningeal whole mounts. Consistent with the sequencing results, we observed a significant increase in collagen density one week after TBI (Figure 3c,d,e). Furthermore, we were also interested in determining whether the fibroblast population was contributing to the inflammatory response following TBI. Of the significantly upregulated genes identified in fibroblasts post-TBI, many of them were related to components of immune system activation and cytokine signaling (Figure 3f).

**Figure 3.**
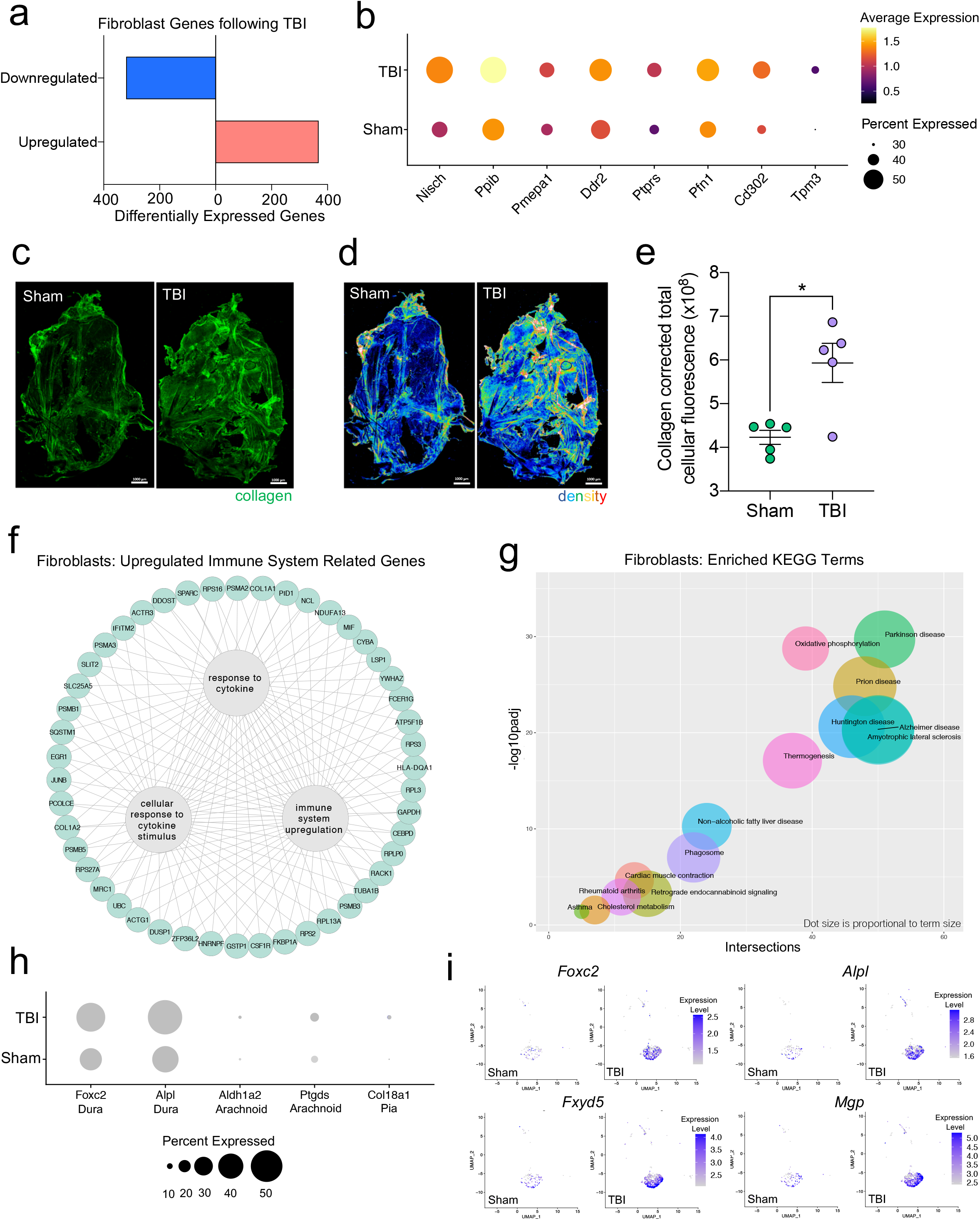
Dural fibroblasts express genes involved in tissue remodeling, cell migration, and immune activation in TBI. Male WT mice at 10 weeks of age received a TBI or Sham procedure. One week later, the meninges from 5 mice per group were harvested, pooled, and processed for scRNA-seq. a) Quantification of the number of upregulated and downregulated fibroblast genes following injury (FDR<0.1). b) Dot plot representation of highlighted fibroblast genes that were significantly upregulated following injury (FDR<0.1). The color and size of each dot represents the average expression and percent of cells expressing each gene, respectively. c-d) Representative images of meningeal whole mounts stained for collagen (green) (c) and a 16 color heatmap of the collagen staining intensity (d), where red is most intense and blue is least intense. e) Quantification of collagen staining intensity using corrected total cellular fluorescence (CTCF). CTCF is calculated as mean fluorescence of meningeal whole mounts - (Area of meningeal whole mount x Mean fluorescence of background). Each data point represents an individual mouse. f) Network map depicting significantly upregulated genes that enriched immune system-related GO terms (FDR<0.1). The lines within the circle indicate which genes contribute to each GO term. g) Scatter plot representation of the top enriched KEGG terms with significantly upregulated genes in the fibroblast population (FDR<0.1). Dot size is proportional to term size. Genes contributing to one KEGG term may also contribute to other KEGG terms. h) Dot plot depicting dural, arachnoid, and pial fibroblasts markers where the size of the circles represents the percent of cells expressing each gene. i) Feature plots of genes characteristic of dural fibroblasts in both Sham and TBI conditions. The color of each data point represents the expression level of the indicated gene within that cell. Graphs were calculated using Seurat by normalizing the dataset, finding the variable features of the dataset, scaling the data, and reducing the dimensionality. Differential gene expression was calculated using the ZINB-WaVE function for zero-enriched datasets and DESeq2. Each data point in a UMAP plot represents a cell. Error bars depict mean ± s.e.m. P values were calculated using a two sample t-test assuming unequal variances. *P<.05. FDR; false discovery rate, p.adj; adjusted p-value.

To explore the cellular and disease pathways that are most affected in fibroblasts post mild head trauma, we identified the KEGG terms enriched by the differentially upregulated genes in the fibroblast group after TBI in comparison to the Sham group. Interestingly, disease pathways related to neurodegenerative diseases, including Parkinson’s disease, Alzheimer’s disease, amyotrophic lateral sclerosis, and prion disease, were some of the most highly upregulated pathways altered in fibroblasts after TBI (Figure 3g). Many of the same terms that contribute to the oxidative phosphorylation KEGG term also contribute to the various disease-related KEGG terms, indicating a change in the metabolic state of the fibroblasts may be underlying disease-related processes.

Given that fibroblasts are present in all three meningeal layers (79), we decided to investigate which layers the fibroblasts inhabited, and which layer was likely responsible for the increase in fibroblasts following TBI. To accomplish this, we examined the expression of molecules that are commonly used to identify the distinct layer of the meninges in which the fibroblast population is likely to reside (79–85). More specifically, recent studies have shown that dural fibroblasts can be identified using the makers *Alpl* and *Foxc2* (80–82), whereas *Ptgds* and *Ald1a2* are unique markers of arachnoid fibroblasts (83, 84) and *Col18a1* is specific to pial fibroblasts (79, 85). As expected, we found that a majority of the fibroblasts in the meninges were from the dura, the thickest layer of the meninges and the layer that is targeted by the method of dissection used in these studies (32, 86). Fewer pia- or arachnoid-resident fibroblasts were present, as anticipated (Figure 3h). When we looked at the expression level of dural fibroblast genes before and after TBI, we saw that several of the markers (e.g., *Foxc2*, *Fxyd5*) were significantly upregulated following injury, and other dural markers, while not expressed at significantly higher levels, were clearly expressed by a higher proportion of total cells (e.g., *Alpl, Mgp*) (Figure 3i). This indicates that the dural compartment likely undergoes an increase in fibroblast cell frequency one week after brain injury, which is also consistent with the increase in collagen density seen in the meninges 1 week after TBI (Figure 3c,d,e). Overall, we observed that the meningeal fibroblast population is highly responsive to TBI and that they upregulate genes enriched in disease-related pathways, immune system activation, and the wound healing response.

### Transcriptional modulation of meningeal lymphocytes in response to mild TBI

Since we observed shifts in the frequencies of some immune cell populations after TBI (Figure 1d,e), we were interested in determining which genes were differentially expressed in meningeal T and B cells after head injury, especially given recent reports that have identified instrumental roles for meningeal lymphocytes in regulating multiple aspects of neurobiology, behavior, and CNS disease (33, 35–37, 39). We independently combined the two T cell populations (“Activated T Cells” and “CD3+ T Cells”) and the B cell populations (“B Cells 1”, “B Cells 2”) to assess differential gene expression. Overall, 151 genes were upregulated and 286 were downregulated following injury in the T cell population, 102 genes were upregulated and 158 were downregulated following injury in the B cell population, and 183 genes were upregulated and 149 were downregulated following injury in the dendritic cell population (Figure 4a). Some of the smaller populations such as NK cells, neutrophils, and plasmacytoid dendritic cells exhibited few to no differentially regulated genes, likely due to the small number of cells present in each of these populations (Figure 4a).

**Figure 4.**
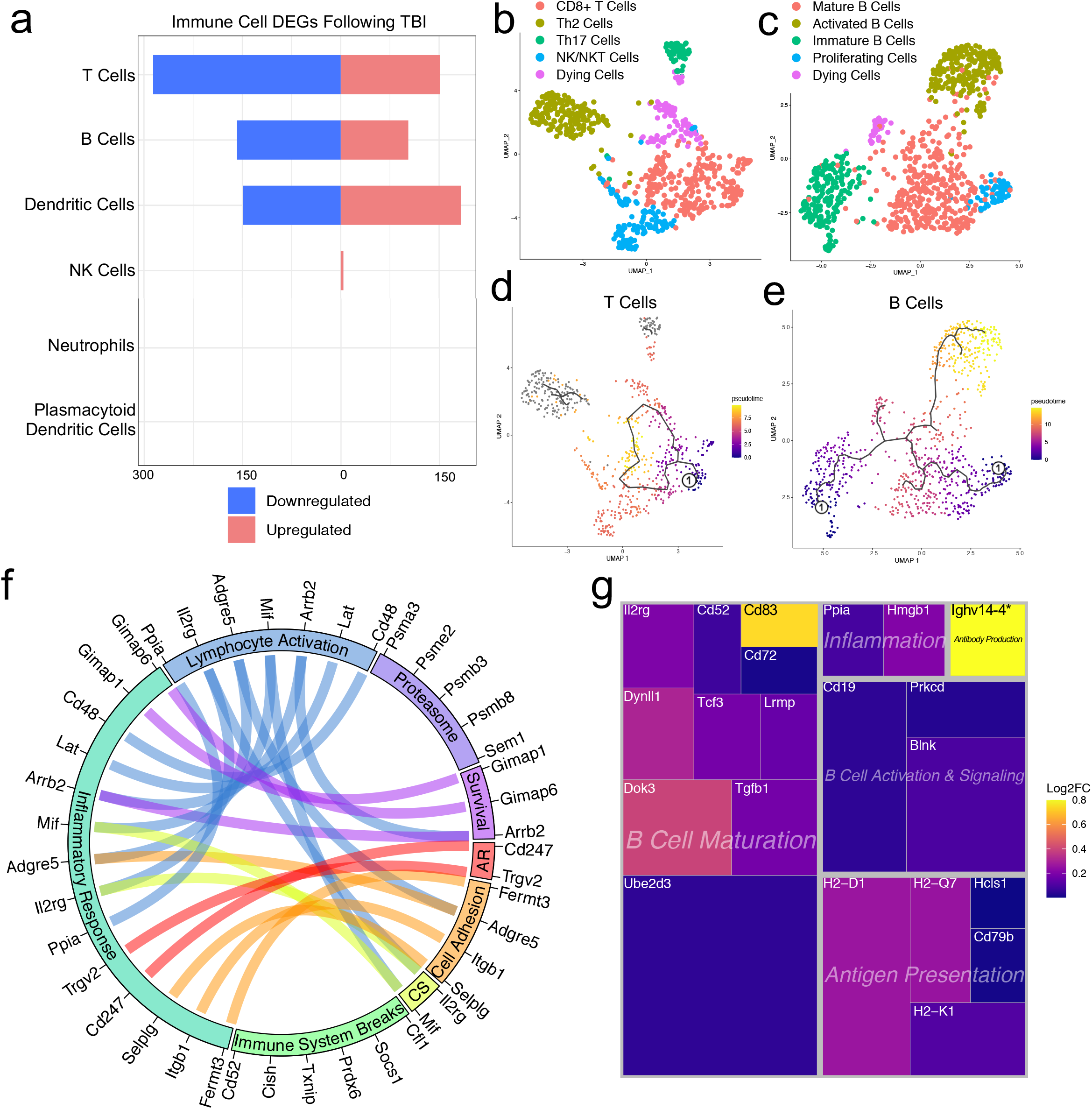
Transcriptional response of meningeal lymphocytes to mild TBI. Male WT mice at 10 weeks of age received a TBI or Sham procedure. One week later, the meninges from 5 mice per group were harvested, pooled, and processed for scRNA-seq. a) Quantification of the number of upregulated and downregulated genes in different immune cell populations following injury (FDR<0.1). b-c) UMAP representation showing re-clustering of the (b) T cell and (c) B cell populations present within the meninges. d-e) UMAP representation of pseudotime cellular trajectory profiles showing (d) T cell and (e) B cell maturation trajectories. The circle with the number “1” represents the root node. The color of each data point represents advancement in pseudotime, with dark purple representing “early” pseudotime and yellow representing “late” pseudotime. The line represents the “path” of pseudotime with intersections representing possible different differentiation events. Grey data points represent cell populations that were not connected in pseudotime with the selected node. f) Circos plot depicting differentially expressed genes in the T cell populations within the TBI meninges (FDR<0.1) associated with different cellular processes. The proportion of the circle’s circumference allocated to each cellular process represents the number of T cell genes associated with that process that are differentially expressed in the TBI meninges. The lines connecting genes within the circle indicate which genes were shared amongst cellular processes. Colors were randomly assigned. g) Treemap depicting significantly upregulated genes in the B cell population and the cellular process to which each gene contributes. The size of the square around each gene represents the Wald statistic, which is used to calculate the overall significance of the change in gene expression (a larger square indicates a larger Wald statistic, which leads to a lower adjusted p-value). The color of the boxes represents log2FC, where purple represents a lower log2FC and yellow represents a higher log2FC. An asterisk (*) indicates that the log2FC of the gene was higher than the scale (*Ighv14-4* had a log2FC of 18.08). Graphs were calculated using Seurat by normalizing the dataset, finding the variable features of the dataset, scaling the data, and reducing the dimensionality. Each data point in a UMAP plot represents a cell. Differential gene expression was calculated using the ZINB-WaVE function for zero-enriched datasets and DESeq2. Pseudotime graphs were created using Monocle. AR, antigen recognition; CS, cytokine signaling; FDR; false discovery rate, log2FC; log 2 fold change.

We were next interested in determining which different T and B cell subsets were present within the meninges, so we re-clustered the cells within these two populations (Figure 4b,c). We found that within the T cell subsets, there was a clear CD8+ T cell population and two T helper cell populations: Th2 cells and Th17 cells (Figure 4b). The Th2 cell sub-cluster expressed highly-significant cluster-defining markers including *Il1rl1* and *Gata3,* which are characteristic of the Th2 subset (87) (Figure 4b, Supplementary Figure 5a,b). Alternatively, the Th17 sub-cluster expressed characteristic markers such as *Il23r, Il17re*, and *Rorc* (88) (Figure 4b, Supplementary Figure 5a,c). The fourth sub-cluster of T cells appears to be comprised of NKT and NK cells, as this population expressed high levels of common NK markers, including *Klrb1c, Ncr1, Klrd1*, and *Klrk1,* and some of these same cells also expressed components of the CD3 co-receptor (*Cd3d, Cd3d,* and *Cd3g*) (Figure 4b, Supplementary Figure 5a,d). The final population represents cells that are likely dying T cells, based on their high expression of mitochondrial genes and *Malat1* (Figure 4b).

Re-clustering of the B cell populations revealed 5 sub-clusters (Figure 4c). One sub-cluster appeared to be comprised of mature B cells given its high expression of B cell maturity marker *Cd37* and the B cell receptor components (*Cd79a* and *Cd79b*). (Supplementary Figure 6a) (89). A second cluster, deemed “Activated B Cells”, was characterized by significant expression of HLA-related genes including *H2-Aa, H2-Eb1,* and *H2-Ab1*, and survival-related genes including *Gimap3, Gimap4*, and *Gimap6*. These activated B cells also highly expressed genes important for adhesion, including *Sell*, which encodes for L-selectin and is a marker for mature B cells (90) (Supplementary Figure 6a,b). A third cluster appeared to be differentiating or immature B cells based on their high expression of *Rag1* and *Rag2* (Supplementary Figure 6a,c). A fourth cluster, deemed “Proliferating Cells” expressed high levels of *Myc* and *Ccnd2* amongst other cell cycle related genes (Supplementary Figure 6a,d). The final population represents cells that are likely dying B cells due to their high expression of *Malat1* (Figure 4c).

In order to determine T and B cell maturation trajectory within the meninges, we performed pseudotemporal analyses using Monocle3 (61). The T cell populations did not demonstrate a strong trajectory in their differentiation status, which is expected given that the populations we identified (Th2, Th17, CD8+ T cells) are all relatively advanced within T cell maturation (Figure 4d). However, when we examined the pseudotemporal trajectory of the B cells, we observed a path that confirmed our initial cluster assignments (Figure 4e). We observed that the B cells earliest in the differentiation trajectory, as demonstrated by the lowest values on the pseudotime scale, were the “Immature B Cells” and “Proliferating Cells” populations, whereas the “Activated B Cells”, that are likely producing antibodies, and “Mature B cells” were the furthest along in the differentiation trajectory (Figure 4e).

Next, we were interested in looking more closely at some of the genes that were significantly upregulated in both the T and B cell populations to determine how these adaptive immune populations were affected following injury. The T cell populations upregulated many genes important for survival (*Gimap1, Gimap6*), activation (*Arrb2, Ppia, Cd48*), cytokine signaling (*Mif, Il2rg*), and antigen recognition (*Cd247*), all of which contributed to an overall increase in inflammatory response gene expression (Figure 4f). Concomitantly, the T cells also upregulated various genes that are known to be involved in the dampening of immune responses such as *Socs1* (Suppressor of Cytokine Signaling-1) and *Cd52* (Figure 4f) (91, 92).

Investigating the genes that were upregulated in the B cell populations following injury, we found that many of these genes fell into the category of “B Cell Maturation”, including *Cd83, Ube2d3*, and *Doc3* (Figure 4g). Other upregulated genes included those important for B cell activation and signaling (*Blnk, Cd19*), antigen presentation (*Cd79b, H2-D1*), and inflammation (*Ppia, Hmgb1*) (Figure 4g). The upregulation of these genes suggests that TBI drives the activation and maturation of B cell populations in the meningeal compartment. Overall, these data demonstrate that both T and B cells upregulate genes involved in activation of adaptive immune responses following head trauma. This upregulation seems to be controlled, as multiple regulatory genes are also simultaneously activated.

### Prominent effects of aging on the meningeal transcriptome that can be further modulated by mild TBI

Given the considerable brain injury-induced alterations in the meningeal transcriptional and cellular landscape that we observed in young mice, we were next interested in investigating whether these changes were preserved or altered with age. It has previously been suggested that inability to properly resolve inflammatory responses in the brain post head trauma contributes to the aggravated disease course commonly seen in the elderly. Therefore, we were also particularly interested to explore potential differences in the resolution of meningeal immune responses following TBI between young and aged mice. To this end, we performed bulk RNA sequencing on the meningeal tissue at 1.5 months post TBI or Sham treatment in both young (10 weeks of age) and aged (20 months of age) mice (Figure 5a), as we predicted that the meningeal injury would have largely resolved 1.5 months post-TBI in young mice. Principal component analysis (PCA) revealed that age was the main driver of differential gene expression, as young and aged groups clustered furthest apart (Figure 5b). However, while the young mice that had received either Sham or TBI clustered together in the PCA plot, the aged Sham and TBI mice clustered further apart, indicating that the effects from TBI may have more long-lasting effects on gene expression in aged mice when compared to young mice (Figure 5b). Indeed, when we looked at the number of differentially expressed genes between all four experimental groups, we saw that there were only a total of 22 differentially expressed genes when comparing Young Sham with Young TBI, while there were a total of 364 differentially expressed genes when comparing Aged Sham with Aged TBI (Figure 5c,d). Interestingly, 1772 differentially expressed genes were identified when comparing Young Sham mice with Aged Sham mice, and 2936 differentially regulated genes were identified when comparing Young TBI mice with Aged TBI mice (Figure 5c,d). This indicates that aging profoundly affects meningeal gene expression and that mild head trauma in aging results in even larger changes in gene expression (Figure 5c,d). Moreover, while the young mice exhibit very few gene expression changes 1.5 months following TBI, the aged mice experience many more alterations in gene expression that last for a longer period of time, which suggests that recovery post-TBI may be delayed with aging.

**Figure 5.**
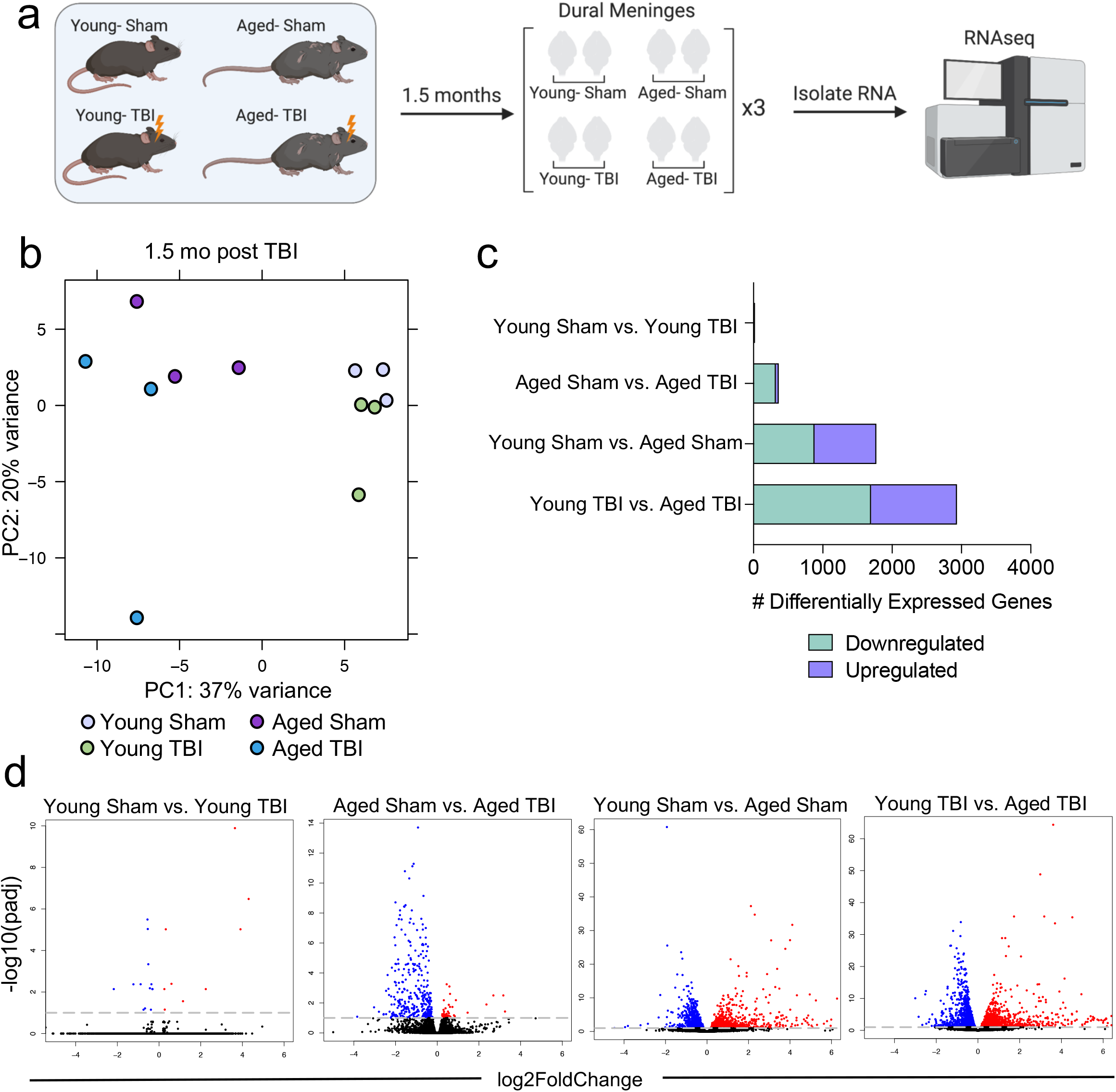
Effects of aging and mild TBI on the meningeal transcriptome. a) Schematic depicting experimental layout. Male WT mice at 10 weeks of age or 20 months of age received a TBI or Sham procedure. 1.5 months later, bulk RNA-seq was performed on the 4 experimental groups with 3 biological replicates per group (each biological replicate consisted of meningeal RNA samples from 2-3 independent mice). b) Principal component analysis (PCA) showing clustering of samples. c) Graphical representation of the upregulated and downregulated genes in all four experimental groups 1.5 month post TBI. d) Volcano plots illustrate the number of differentially expressed genes with statistically significant differences denoted in blue and red (FDR<0.1). Blue data points represent significantly downregulated genes and red data points represent significantly upregulated genes. FDR and p-values were calculated with DESeq2 using the Wald test for significance following fitting to a negative binomial linear model and the Benjamini-Hochberg procedure to control for false discoveries. FDR; false discovery rate.

Because aging itself resulted in substantially different gene expression patterns, we decided to look more closely at these differences. Upon examining the top 20 upregulated and downregulated genes in the Aged Sham mice as compared to the Young Sham mice, we noticed a striking upregulation in genes important for antibody production by B cells (Figure 6a). In fact, one half of the top 20 upregulated genes fell into this category (Figure 6a). When we examined the top GO biological processes that were enriched by the significantly activated genes in the Young Sham mice versus Aged Sham mice comparison, we saw that immune and defense responses were among the most highly upregulated (Figure 6b), indicating that the cells within the aged meninges have grossly upregulated their immune response, even in homeostatic conditions.

**Figure 6.**
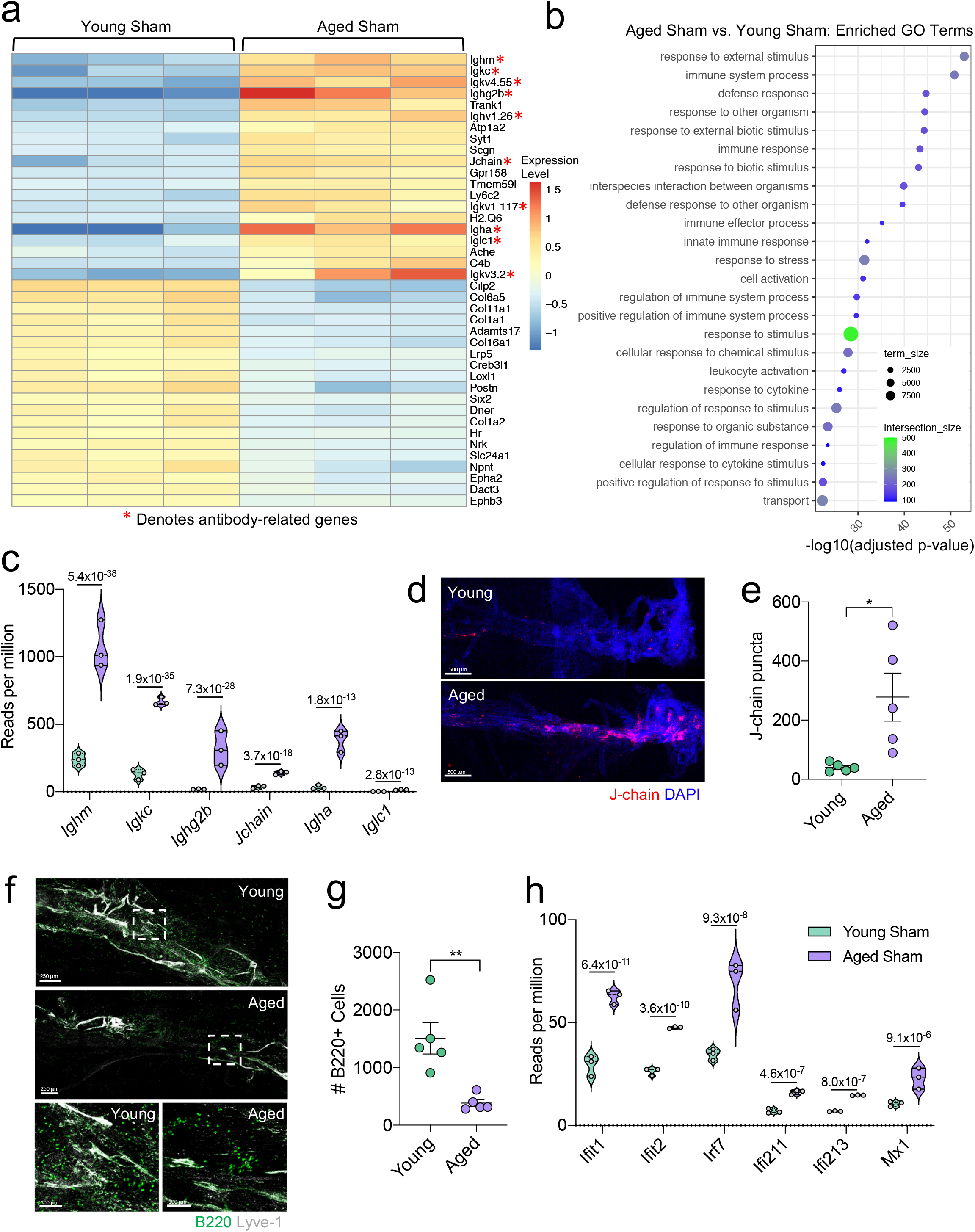
Aging promotes the upregulation of meningeal genes involved in type I IFN and antibody signaling. Male WT mice at 10 weeks of age or 20 months of age received a TBI or Sham procedure. 1.5 months later, bulk RNA-seq was performed on the 4 experimental groups with 3 biological replicates per group (each biological replicate consisted of meningeal RNA samples from 2-3 independent mice). a) Heatmap representation of the top 20 most significantly upregulated and downregulated (FDR<0.1) genes in the Young Sham vs. Aged Sham groups. The red asterisk (*) indicates genes associated with antibody production. b) Dot plot of GO term biological processes shows enrichment of immune-related pathways with differentially expressed genes between young mice as compared to aged mice. Color and size of each dot represent the size of the GO term and the number of upregulated genes that contribute to each term, respectively. c) Violin plot depicting counts of significantly activated antibody and B cell related genes in response to age (FDR<0.1). The number above each comparison on the graph represents the adjusted p-value calculated for each gene using DESeq2. The central line within each plot represents the median of the data set. The upper and lower boundaries of the box represent the third (Q3) and first (Q1) quartiles respectively. The violin plot encompasses the three biological replicates. The width of the violin plot represents the frequency of observations at that given y-value. Therefore, the wider the violin plot, the higher the frequency of observations. The meninges of 5 young Sham mice and 5 aged Sham mice were harvested for each immunohistochemical experiment. (d) Representative images from a young Sham mouse and aged Sham mouse showing a region of the SSS stained with J-chain (red) and Lyve-1 (grey) (e) and quantification of J-chain puncta in meningeal whole mounts along the SSS. f) Representative images of the transverse sinus in young and aged mice stained with B220 (green) and Lyve-1 (grey). The dashed box on the top two images corresponds to the higher magnification images depicted below. g) Quantification of the number of B220 cells along the entire transverse sinus. h) Violin plot depicting counts of significantly activated type-I interferon related genes in response to age (FDR<0.1). The violin plot parameters are the same as describe for (c). FDR and p-values in (a-c,i) were calculated with DESeq2 using the Wald test for significance following fitting to a negative binomial linear model and the Benjamini-Hochberg procedure to control for false discoveries. Error bars in (e,g) depict mean ± s.e.m. P values in (e,g) were calculated using an unpaired two-sample t-test assuming unequal variances. *P<.05, **P<0.01. FDR; false discovery rate.

Due to the striking nature of the upregulation of antibody production-related genes, and recent findings that report an increase in IgA-secreting plasma cells with age (93), we more closely examined some of these genes (Figure 6c). We found highly significant upregulations in genes related to the immunoglobulin heavy chain (*Ighm, Ighg2b, Igha*), light chain (*Igkc*), and components of IgA or IgM antibodies (*Jchain*) (Figure 6c). Using immunohistochemistry on meningeal whole mounts, we confirmed that aged meninges have significantly increased J chain deposition that is concentrated along the sinuses (Figure 6d,e). Next, we wanted to determine whether the increased antibody-related gene production we saw in aged mice was due to an overall increased number of B cells. Interestingly, we saw that the number cells expressing B220 along the meningeal transverse sinus in mice was significantly lower in aged mice (Figure 6f,g), which is in contrast to other recent studies have shown that B cells comprise a larger proportion and number of cells in aged dural meninges (94, 95). Differences in our data in compared to this published data may reflect regional differences in B cell populations along the transverse sinus given the impaired meningeal lymphatics seen in aged mice. Furthermore, B cells may comprise an overall greater proportion of cells in aged meninges, but given the overall decreased cellularity in aged meninges, may represent a smaller absolute number when compared with young meninges. These data show aged meninges harbor mature B cells that have a higher expression of antibody production-related genes. Overall, this suggests that the composition of the B cell population in aged mice may be substantially different than in young mice; however, future studies are needed to formally evaluate this in greater detail.

In addition to the antibody-related gene upregulation, we also observed increased expression of type I interferon (IFN)-related genes (Figure 6h). Type I IFNs have been shown to be upregulated in the brain parenchyma in various neurological disorders, where they are generally thought to play deleterious roles in promoting disease pathogenesis (70, 71, 96–98). Our data indicate that this type I IFN signature is also seen within the meningeal compartment of aged mice. Amongst others, we saw highly significant increases in type I IFN related genes including *Ifit1, Ifit2, Irf7, Ifi213,* and *Mx1* (Figure 6h). These findings demonstrate that aging promotes profound alterations in the meningeal transcriptome. Moreover, the upregulation of antibody genes and type I IFN related-genes suggests an overall elevation in immune activation status in the aged meninges.

### Injury in aged mice results in prolonged inflammatory responses

In order to assess the unique transcriptional response to TBI in aged compared to young mice, we analyzed the transcriptional response that is exclusive to the Young TBI vs Aged TBI comparison and not shared with the Young Sham vs Aged Sham comparison. Of the differentially expressed genes, there were 1186 genes that were shared amongst these two comparisons, 1750 genes that were unique to the Young TBI vs Aged TBI comparison and 586 genes that were unique to the Young Sham vs Aged Sham comparison (Figure 7a). While aging and TBI each individually lead to changes in gene expression which have some overlap, the double hit of TBI with old age was found to induce an even larger change in gene expression than either condition alone.

**Figure 7.**
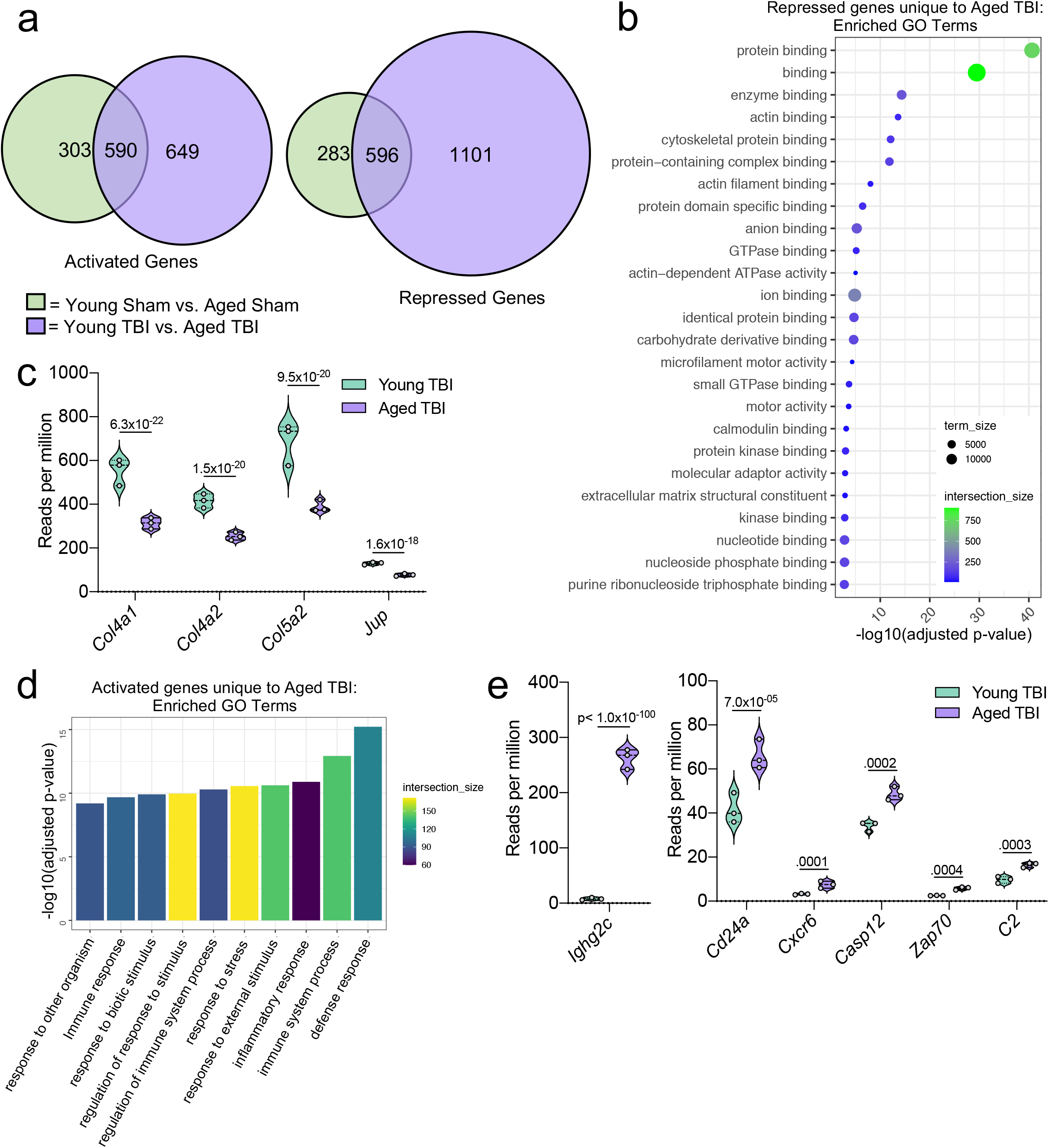
Aging and mild TBI together promote a unique meningeal transcriptional signature. Male WT mice at 10 weeks of age or 20 months of age received a TBI or Sham procedure. 1.5 months later, bulk RNA-seq was performed on the 4 experimental groups with 3 biological replicates per group (each biological replicate consisted of meningeal RNA samples from 2-3 independent mice). a) Venn diagram depicting unique and shared differentially regulated genes between the Young Sham vs Aged Sham and Young TBI vs Aged TBI groups (FDR<0.1). Circle size is roughly correlated with gene number. b) Dot plot showing GO term molecular functions enriched by the repressed genes unique to the Young TBI vs Aged TBI comparison. The color and size of each dot represents the size of the GO term and the number of upregulated genes that contribute to each term, respectively. c) Violin plots depicting counts of significantly repressed extracellular matrix related genes (FDR<0.1). d) Bar plot shows enrichment of GO term biological processes related to the immune system with the genes unique to the Young TBI vs Aged TBI comparison. The color of each bar represents the number of upregulated genes that contribute to each GO term. e) Violin plots depicting counts of significantly activated immune-related genes (FDR<0.1). (c,e) Each statistic represents the adjusted p-value calculated for each gene using DESeq2. The central line within each plot represents the median of the data set. The upper and lower boundaries of the box represent the third (Q3) and first (Q1) quartiles respectively. The violin plot encompasses the three biological repeats. The width of the violin plot represents the frequency of observations at that given y-value. Therefore, the wider the violin plot, the higher the frequency of observations. FDR and p-values were calculated with DESeq2 using the Wald test for significance following fitting to a negative binomial linear model and the Benjamini-Hochberg procedure to control for false discoveries.

Next, we more closely examined the 1750 differentially expressed genes unique to the Young TBI vs Aged TBI group that were not shared with the Young Sham vs Aged Sham comparison (Figure 7a). Using the GO molecular function terms, we saw that of the 1101 repressed genes, many of these genes are involved in binding processes, including protein binding and cytoskeletal binding (Figure 7b). When we looked more closely at the top repressed genes unique to the Aged TBI versus Young TBI comparison, we observed that many of these genes encode for collagenases (*Col4a1, Col4a2,* and *Col5a2*) and other molecules involved in regulating cellular junctions (*Jup*) (Figure 7c). These pathways likely aid in the wound healing response of the meninges but are downregulated after brain injury in aging.

Additionally, we looked into the genes that were uniquely activated in the Aged TBI mice as compared to the Young TBI mice. We found that genes associated with immune activation were profoundly upregulated in aged TBI mice in comparison to their young TBI counterparts (Figure 7d). The most enriched GO biological processes included the “defense response” and “immune system process” (Figure 7d). Some of the genes that contributed to the upregulation of these immune-related terms included those associated with immunoglobulin production (*Ighg2c*), T and B cell signaling (*Cd24a, Zap70, Cxcr6*), and cell death (*Casp12, C2*) (Figure 7e). In summary, these findings highlight some of the distinct changes seen in the aged meningeal tissue following TBI. Specifically, we find that mild TBI in aged mice results in prolonged activation of immune genes and decreased expression of genes involved in extracellular matrix remodeling and the maintenance of cellular junctions. Furthermore, we report that while the meningeal transcriptome in young mice returns almost completely to baseline resting levels by 1.5 months post mild head injury, the aged meninges, in contrast, continue to exhibit substantial and protracted transcriptional alterations related to head injury.

## DISCUSSION

Findings from these studies highlight the heterogeneous and dynamic nature of the meninges in response to TBI and aging. Following TBI in young mice, there is an enrichment of fibroblast and macrophage populations in the meninges, as well as an upregulation in genes associated with immune activation. Interestingly, the gene expression patterns of the meninges are drastically altered in aging, with large upregulations in genes involved in immunoglobulin production and type I IFN signaling. Upon injury, the aged meninges downregulate the production of genes related to collagenase production and other genes important for extracellular matrix maintenance and cell junction formation, while they continue to upregulate genes involved in immune signaling. Moreover, the aged meninges experience a much more prolonged and substantial response to injury than the meninges in young mice, which have largely returned to baseline by 1.5 months post-injury.

While our overall knowledge of meningeal biology in TBI remains limited, recent work has begun to uncover roles for meningeal macrophages in head trauma (27, 28). For instance, studies by Roth et al. demonstrated that the release of reactive oxygen species (ROS), which occurs as part of meningeal macrophage cell death, can trigger subsequent tissue damage in the brain parenchyma (28). Interestingly, they showed that blocking ROS release by dying meningeal macrophages is also effective in minimizing cell death in the brain parenchyma. Collectively, these results suggest that the meningeal response to TBI directly affects cells in the brain as well (28). More recently, this same group also demonstrated that meningeal macrophages play important roles in coordinating meningeal remodeling and vascular repair following mild head trauma (27). Here, they showed that distinct macrophage populations function within defined regions in and around the injury site. For example, they identified that inflammatory myelomonocytic cells work in the core of the lesion where dead cells are abundant, whereas wound-healing macrophages are present along the perimeter of the injury where they work to restore blood vasculature and clear fibrin (27).

Recent studies have also begun to uncover critical roles for meningeal lymphocytes in various neurological disorders as well as in the regulation of basic neurological functions and behavior (32, 33, 35–37, 86, 94, 95, 99). Yet, surprisingly little is currently known with regard to how head trauma impacts adaptive immunity in the meninges. Here, we show that adaptive immune cells found in the meninges of young mice upregulate genes essential for activation and maturation at one week post-TBI. Furthermore, in both aging alone and in aged mice following brain injury, we observed a massive upregulation of genes important for antibody production by B cells. Interestingly, emerging studies have proposed that IgA-producing plasma B cells survey the meninges and that their IgA secretion provides an “immunological barrier” to prevent potential pathogens from gaining access into the brain parenchyma (93). While speculative, perhaps this upregulation in antibody production genes in the meninges is a protective strategy that is mobilized to counteract the loss of blood-brain barrier (BBB) integrity that can occur both in aging and following TBI (51, 100). Nevertheless, further studies are required to follow up on the significance and function of this massive elevation in antibody-related genes that is seen in the meninges as a result of aging and head trauma.

For the studies that appear in this paper we paid special attention to the transcriptional response that occurs in meningeal macrophages, fibroblasts, and lymphocytes given their high abundance in the meninges as well as emerging evidence suggesting important roles for these immune cell lineages in TBI pathogenesis. However, it should be noted that there are many other populations of cells present in the meninges in which we were not able to adequately assess differential gene expression due to their small population sizes. In future studies, it will be important to further interrogate the functions of these less abundant cell populations in TBI. Notably, the presence of plasmacytoid dendritic cells within the meninges is intriguing as they are known to be potent producers of type I IFNs (101). It is possible that plasmacytoid cells could be contributing to the type I IFN signature seen with aging and TBI in the meninges; however, future work is needed to formally investigate this.

It is also known that sex differences play a role in outcomes after TBI both in human and animal models (102). While men have a higher likelihood of sustaining a TBI, women have a higher likelihood of suffering worse outcomes (102). Due to the number of other variables we were already considering for this study (age, time and injury status), we were unable to include sex as a variable for our sequencing data. This leaves further opportunities for investigation into how sex impacts the meningeal transcriptome in the context of both TBI and aging. High throughput sequencing techniques provide unique opportunities to understanding how sex affects CNS tissues at a cellular and transcriptional level.

While the meningeal immune response to head trauma in aged mice appears to be largely upregulated, the meningeal immune response in young mice following injury appears to be held in check. One week following injury, although there is an upregulation in genes important for the inflammatory response in the macrophage and adaptive immune cell populations, there are other upregulated genes that are important for dampening this same immune response and for promoting wound healing. For instance, the subsets of macrophages whose frequencies increase the most following injury are “Anti-Inflammatory” and “Resolution Phase” macrophages, both of which have been reported to exert wound-healing properties in injury models (27, 75). Furthermore, while T cells were observed to upregulate genes associated with immune activation, survival, and adhesion, they also displayed increased expression of genes involved in dampening cytokine signaling and controlling the immune response in the meninges of young mice (e.g., *Cfl1, Socs2*, and *Cd52*). Moreover, the vast majority of these differentially expressed genes seen in young meninges at one week post-injury return to resting levels by 1.5 months following head trauma, suggesting a resolution of inflammation and a restoration of homeostasis in young mice. In contrast, aged mice do not appear to have this same success in resolving inflammatory responses following head trauma. In addition to the baseline inflammatory state of aged meninges that is characterized by increased expression of genes related to type I IFN signaling and antibody production, aged mice that received a TBI were found to further upregulate genes involved in driving inflammatory responses. Moreover, these injured aged mice were also shown to downregulate numerous genes involved in extracellular matrix reorganization and collagen production, which are two processes that are necessary for proper tissue regeneration. These transcriptional alterations in aged meninges persist beyond a month following injury, with no indications of resolution.

It is well known that aged individuals have a higher morbidity and mortality than young individuals when experiencing a similar severity brain injury (11). The explanation for why the elderly experience these poorer outcomes following TBI is likely complex and multifaceted. Many studies have highlighted baseline changes in the aged brain that has been speculated to prime the elderly for differential responses following injury, including changes in the BBB, microglial dysfunction, and an overall increase in neuroinflammation (23, 100, 103). Furthermore, other findings support changes in the response to injury in the aged brain, including alterations in the type and number of immune cells recruited to the injury site, further increases in inflammatory gene signatures in the brain parenchyma, and elevated production of potentially neurotoxic molecules such as ROS and type I IFNs (17, 20, 21, 23–26, 70, 71, 96–98, 104). Our findings indicate that the meninges may also play a role in this differential response to head trauma seen in aging. In particular, it is possible that the increased baseline type I IFN gene signature and antibody production observed in aging potentially renders the aged brain prone to more severe clinical outcomes post-TBI.

Recent studies have implicated the meningeal lymphatic system, which resides in the dura, in modulating inflammation in the brain following TBI and sub-arachnoid hemorrhage (52, 105, 106). In these studies, impairments in the meningeal lymphatic system prior to brain injury were found to result in increased gliosis and worsened behavioral outcomes (52, 105). Interestingly, the meningeal lymphatic system is also known to be impaired in aging (107–109), and we have previously shown that the rejuvenation of the meningeal lymphatic vasculature in aged mice dampens the subsequent gliosis following TBI (52). How the meningeal lymphatic system might modulate meningeal immunity before and after injury remains to be investigated. Furthermore, whether the meningeal lymphatic impairment in aging contributes to the overall increase in inflammation seen in the aged meninges is another area for future investigation.

### Conclusions

Overall, the findings presented here provide new insights into the meningeal response to brain injury and aging. We show that TBI results in broad gene expression changes in discrete cell populations following injury in young mice. Specifically, we demonstrate that there is an increase in the frequency of fibroblasts and macrophages one week following injury in young mice. Furthermore, we provide evidence that the transcriptional environment in the aged meninges is drastically altered. At baseline, the aged meninges show increases in gene expression patterns associated with type I IFN signaling and antibody production by B cells. However, upon injury, the aged meninges further upregulate genes involved in immune system activation, while downregulating genes critical for tissue remodeling. Improved understanding of how the meninges respond to brain injury in youth and aging will help shed light on why the elderly have poor outcomes following TBI and may help to identify opportunities for targeted therapies to improve outcomes following TBI.

## ABBREVIATIONS

AR: antigen recognition
BBB: blood brain barrier
CNS: central nervous system
CS: cytokine signaling
CSF: cerebrospinal fluid
DEG: differentially expressed gene
FDR: false discovery rate
GO: gene ontology
IFN: interferon
ISF: interstitial fluid
MRI: magnetic resonance imaging
p.adj: adjusted p-value
PC: principal component
PCA: principal component analysis
RNA-seq: RNA sequencing
ROS: reactive oxygen species
scRNA-seq: single-cell RNA sequencing
TBI: traumatic brain injury
UMAP: uniform manifold approximation and projection

## DECLARATIONS

### Ethics approval and consent to participate

All mouse experiments were performed in accordance with the relevant guidelines and regulations of the University of Virginia and approved by the University of Virginia Animal Care and Use Committee.

### Consent for publication

All authors have reviewed this manuscript and agreed to publish it in the current form.

### Availability of data and material

All data and genetic material used for this paper are available from the authors on request. All code used for analysis is available at [https://github.com/danielshapiro1/MeningealTransciptome] or upon request.

## Funding

This work was supported by The National Institutes of Health/National Institute of Neurological Disorders and Stroke (R01NS106383; awarded to J.R.L.), The Alzheimer’s Association (AARG-18-566113; awarded to J.R.L.), The Owens Family Foundation (Awarded to J.R.L.), and The University of Virginia Research and Development Award (Awarded to J.R.L.). A.C.B., A.B.D. and W.F.M. were supported by a Medical Scientist Training Program Grant (5T32GM007267-38). A.C.B was supported by an Immunology Training Grant (5T32AI007496-25), a Wagner Fellowship, and the National Institutes on Aging (NIA, F30AG069396-01). A.B.D was supported by a Biomedical Data Sciences Training Grant (T32LM012416).

### Authors’ contributions

A.C.B and J.R.L. designed the study; A.C.B., D.A.S., K.R.B, and A.R.M performed experiments. A.B.D., D.A.S., and W.F.M. contributed to data analysis. A.C.B. and J.R.L. analyzed data and wrote the manuscript; J.R.L. oversaw the project. All authors read and approved the final manuscript.

## Acknowledgements

We thank members of the Lukens lab and the Center for Brain Immunology and Glia (BIG) for valuable discussions. Graphical illustrations in Figure 1, Figure 5, Supplementary Figure 1, and the graphical abstract were made using BioRender (https://biorender.com/).

## SUPPLEMENTARY FIGURE LEGENDS

**Supplementary Figure 1. Initial brain and meningeal response following TBI**. Male C57BL/6J wild-type (WT) mice at 10 weeks of age received a TBI or Sham procedure and then the brains or meninges were harvested for immunohistochemistry at 24 hours. a) Schematic depicting the location of the TBI in relation to dorsal anatomical structures. b) Representative images of meningeal whole mounts stained with DAPI (blue), CD31 (green) and Lyve-1 (grey). c) Quantification of the percent area of CD31 in each meningeal whole mount. Each data point represents an individual mouse. d) Representative images of brains with injury site (right) taken 24 hours following TBI stained with DAPI (blue), Iba1 (green), NeuN (grey) and GFAP (yellow). e) Representative high magnification images (63x) of Iba1+ and GFAP+ cells (microglia/macrophages and reactive astrocytes respectively) and f) quantification of the percent area of GFAP and Iba1 positive cells in the hemisphere ipsilateral to the injury 24 hours after TBI. Each data point represents an individual mouse. g-i) Meningeal whole mounts stained with MHCII (red) taken 24 hours post TBI. Dashed boxes represent zoomed areas of transverse sinus shown in (g). i) Quantification of the % area MHCII staining in each meningeal whole mount. Each dot represents one mouse. Error bars depict mean ± s.e.m. P values were calculated using the students t-test. *P<.05.

**Supplementary Figure 2. Cluster-defining genes for single cell populations**. Male WT mice at 10 weeks of age received a TBI or Sham procedure. One week later, the meninges from 5 mice per group were harvested, pooled, and processed for scRNA-seq. Tables depicting the top 20 most significant cluster-defining genes for clusters 1-21, which were produced by normalizing the dataset, finding the variable features of the dataset, scaling the data, and reducing the dimensionality. Each gene is displayed with its corresponding P_adj. P_adj, adjusted p-value.

**Supplementary Figure 3. Stress and processing related genes after single cell RNA-sequencing.** Male WT mice at 10 weeks of age received a TBI or Sham procedure. One week later, the meninges from 5 mice per group were harvested, pooled, and processed for scRNA-seq. Violin plots depicting various genes split by experimental group: Sham (sage) and TBI (purple). Each dot represents an individual cell. The width of the violin plot represents the frequency of observations at that given y-value. Therefore, the wider the violin plot, the higher the frequency of observations. Plots without sage or purple coloring did not have enough cells expressing the gene to create the plot. Graphs were calculated using Seurat by normalizing the dataset, finding the variable features of the dataset, scaling the data, and reducing the dimensionality.

**Supplementary Figure 4. Cluster-defining genes for macrophage subpopulations**. Male WT mice at 10 weeks of age received a TBI or Sham procedure. One week later, the meninges from 5 mice per group were harvested, pooled, and processed for scRNA-seq. a) Tables depicting the top 10 most significant cluster-defining genes for the five identified macrophage populations. b-e) Feature plots showing expression patterns of (b) Anti-Inflammatory, (c) Resolution Phase, (d) Inflammatory 1, and (e) Inflammatory 2 macrophage cluster-defining genes. The color of each data point represents the expression level of the indicated gene within that cell. Graphs were calculated using Seurat by normalizing the dataset, finding the variable features of the dataset, scaling the data, and reducing the dimensionality. P_adj, adjusted p-value.

**Supplementary Figure 5. Cluster-defining genes for T cell subpopulations**. Male WT mice at 10 weeks of age received a TBI or Sham procedure. One week later, the meninges from 5 mice per group were harvested, pooled, and processed for scRNA-seq. a) Tables depicting the top 10 most significant cluster-defining genes for the four identified T cell populations. b-d) Feature plots showing expression patterns of (b) Th2, (c) Th17 and (d) NK/NKT T cell subset cluster-defining genes. The color of each data point represents the expression level of the indicated gene within that cell. Graphs were calculated using Seurat by normalizing the dataset, finding the variable features of the dataset, scaling the data, and reducing the dimensionality. P_adj, adjusted p-value.

**Supplementary Figure 6. Cluster-defining genes for B cell subpopulations**. Male WT mice at 10 weeks of age received a TBI or Sham procedure. One week later, the meninges from 5 mice per group were harvested, pooled, and processed for scRNA-seq. a) Tables depicting the top 10 most significant cluster-defining genes for the four identified B cell populations. b-d) Feature plots showing expression patterns of (b) Activated, (c) Immature and (d) Proliferating B cell subset cluster-defining genes. The color of each data point represents the expression level of the indicated gene within that cell. Graphs were calculated using Seurat by normalizing the dataset, finding the variable features of the dataset, scaling the data, and, reducing the dimensionality. P_adj, adjusted p-value.

## LITERATURE CITED

1. Marin JR, Weaver MD, Mannix RC. Burden of USA hospital charges for traumatic brain injury. Brain Inj. 2017;31(1):24–31.

2. Roozenbeek B, Maas AI, Menon DK. Changing patterns in the epidemiology of traumatic brain injury. Nat Rev Neurol. 2013;9(4):231–6.

3. Smith DH, Johnson VE, Stewart W. Chronic neuropathologies of single and repetitive TBI: substrates of dementia? Nat Rev Neurol. 2013;9(4):211–21.

4. McKee AC, Cantu RC, Nowinski CJ, Hedley-Whyte ET, Gavett BE, Budson AE, et al. Chronic traumatic encephalopathy in athletes: progressive tauopathy after repetitive head injury. J Neuropathol Exp Neurol. 2009;68(7):709–35.

5. Mark Faul VC. Epidemiology of traumatic brain injury. Handbook of Clinical Neurology. 2015;127:3–13.

6. Selassie AW, Zaloshnja E, Langlois JA, Miller T, Jones P, Steiner C. Incidence of long-term disability following traumatic brain injury hospitalization, United States, 2003. J Head Trauma Rehabil. 2008;23(2):123–31.

7. Kang JH, Lin HC. Increased risk of multiple sclerosis after traumatic brain injury: a nationwide population-based study. J Neurotrauma. 2012;29(1):90–5.

8. Jesse R Fann ARR, Henrik Schou PEdersen, Morten Fenger-Gron, Jakob Christensen, Michael Eriksen Benros, Mogens Vestergaard. Long-Term risk of demetia among people with traumatic brain injury in Denmark: a population-based observational cohort study. The Lancet Psychiatry. 2018;5(5):424–31.

9. Frost RB, Farrer TJ, Primosch M, Hedges DW. Prevalence of traumatic brain injury in the general adult population: a meta-analysis. Neuroepidemiology. 2013;40(3):154–9.

10. Centers for Disease Control and Prevention USDoHaHS. Surveillance Report of Traumatic Brain Injury-related Emergency Department Visits, Hospitalizations, and Deaths—United Sates, 2014. 2019.

11. Susman M, DiRusso S, Sullivan T, Risucci D, Nealon P, Cuff S, et al. Traumatic Brain Injury in the Elderly: Increased Mortality and Worse Functional Outcomes at Discharge Despite Lower Injury Severity. J Trauma. 2002;53(2):219–23.

12. Erturk A, Mentz S, Stout EE, Hedehus M, Dominguez SL, Neumaier L, et al. Interfering with the Chronic Immune Response Rescues Chronic Degeneration After Traumatic Brain Injury. J Neurosci. 2016;36(38):9962–75.

13. Winston CN, Noel A, Neustadtl A, Parsadanian M, Barton DJ, Chellappa D, et al. Dendritic Spine Loss and Chronic White Matter Inflammation in a Mouse Model of Highly Repetitive Head Trauma. Am J Pathol. 2016;186(3):552–67.

14. Corps KN, Roth TL, McGavern DB. Inflammation and neuroprotection in traumatic brain injury. JAMA Neurol. 2015;72(3):355–62.

15. Johnson VE, Stewart JE, Begbie FD, Trojanowski JQ, Smith DH, Stewart W. Inflammation and white matter degeneration persist for years after a single traumatic brain injury. Brain. 2013;136(Pt 1):28–42.

16. Schimmel SJ, Acosta S, Lozano D. Neuroinflammation in traumatic brain injury: A chronic response to an acute injury. Brain Circ. 2017;3(3):135–42.

17. Chou A, Krukowski K, Morganti JM, Riparip LK, Rosi S. Persistent Infiltration and Impaired Response of Peripherally-Derived Monocytes after Traumatic Brain Injury in the Aged Brain. Int J Mol Sci. 2018;19(6).

18. McKee CA, Lukens JR. Emerging Roles for the Immune System in Traumatic Brain Injury. Front Immunol. 2016;7:556.

19. Witcher KG, Bray CE, Chunchai T, Zhao F, O’Neil SM, Gordillo AJ, et al. Traumatic brain injury causes chronic cortical inflammation and neuronal dysfunction mediated by microglia. J Neurosci. 2021.

20. Morganti JM, Riparip LK, Chou A, Liu S, Gupta N, Rosi S. Age exacerbates the CCR2/5-mediated neuroinflammatory response to traumatic brain injury. J Neuroinflammation. 2016;13(1):80.

21. Ritzel RM, Lai YJ, Crapser JD, Patel AR, Schrecengost A, Grenier JM, et al. Aging alters the immunological response to ischemic stroke. Acta Neuropathol. 2018;136(1):89–110.

22. Webster SJ, Van Eldik LJ, Watterson DM, Bachstetter AD. Closed head injury in an age-related Alzheimer mouse model leads to an altered neuroinflammatory response and persistent cognitive impairment. J Neurosci. 2015;35(16):6554–69.

23. Androvic P, Kirdajova D, Tureckova J, Zucha D, Rohlova E, Abaffy P, et al. Decoding the Transcriptional Response to Ischemic Stroke in Young and Aged Mouse Brain. Cell Rep. 2020;31(11):107777.

24. Ritzel RM, Doran SJ, Glaser EP, Meadows VE, Faden AI, Stoica BA, et al. Old age increases microglial senescence, exacerbates secondary neuroinflammation, and worsens neurological outcomes after acute traumatic brain injury in mice. Neurobiol Aging. 2019;77:194–206.

25. Kumar A, Stoica BA, Sabirzhanov B, Burns MP, Faden AI, Loane DJ. Traumatic brain injury in aged animals increases lesion size and chronically alters microglial/macrophage classical and alternative activation states. Neurobiol Aging. 2013;34(5):1397–411.

26. Krukowski K, Chou A, Feng X, Tiret B, Paladini MS, Riparip LK, et al. Traumatic Brain Injury in Aged Mice Induces Chronic Microglia Activation, Synapse Loss, and Complement-Dependent Memory Deficits. Int J Mol Sci. 2018;19(12).

27. Russo MV, Latour LL, McGavern DB. Distinct myeloid cell subsets promote meningeal remodeling and vascular repair after mild traumatic brain injury. Nat Immunol. 2018;19(5):442–52.

28. Roth TL, Nayak D, Atanasijevic T, Koretsky AP, Latour LL, McGavern DB. Transcranial amelioration of inflammation and cell death after brain injury. Nature. 2014;505(7482):223–8.

29. Turtzo LC, Jikaria N, Cota MR, Williford JP, Uche V, Davis T, et al. Meningeal blood-brain barrier disruption in acute traumatic brain injury. Brain Communications. 2020.

30. Aspelund A, Antila S, Proulx ST, Karlsen TV, Karaman S, Detmar M, et al. A dural lymphatic vascular system that drains brain interstitial fluid and macromolecules. J Exp Med. 2015;212(7):991–9.

31. Louveau A, Smirnov I, Keyes TJ, Eccles JD, Rouhani SJ, Peske JD, et al. Structural and functional features of central nervous system lymphatic vessels. Nature. 2015;523(7560):337–41.

32. Alves de Lima K, Rustenhoven J, Kipnis J. Meningeal Immunity and Its Function in Maintenance of the Central Nervous System in Health and Disease. Annu Rev Immunol. 2020;38:597–620.

33. Alves de Lima K, Rustenhoven J, Da Mesquita S, Wall M, Salvador AF, Smirnov I, et al. Meningeal γδ T cells regulate anxiety-like behavior via IL-17a signaling in neurons. Nature Immunology. 2020;21(11):1421–9.

34. Rustenhoven J, Drieu A, Mamuladze T, de Lima KA, Dykstra T, Wall M, et al. Functional characterization of the dural sinuses as a neuroimmune interface. Cell. 2021.

35. Filiano AJ, Xu Y, Tustison NJ, Marsh RL, Baker W, Smirnov I, et al. Unexpected role of interferon-gamma in regulating neuronal connectivity and social behaviour. Nature. 2016;535(7612):425–9.

36. Derecki NC, Cardani AN, Yang CH, Quinnies KM, Crihfield A, Lynch KR, et al. Regulation of learning and memory by meningeal immunity: a key role for IL-4. J Exp Med. 2010;207(5):1067–80.

37. Ribeiro M, Brigas HC, Temido-Ferreira M, Pousinha PA, Regen T, Santa C, et al. Meningeal gammadelta T cell-derived IL-17 controls synaptic plasticity and short-term memory. Sci Immunol. 2019;4(40).

38. Da Mesquita S, Herz J, Wall M, Dykstra T, de Lima KA, Norris GT, et al. Aging-associated deficit in CCR7 is linked to worsened glymphatic function, cognition, neuroinflammation, and beta-amyloid pathology. Sci Adv. 2021;7(21).

39. Gate D, Saligrama N, Leventhal O, Yang AC, Unger MS, Middeldorp J, et al. Clonally expanded CD8 T cells patrol the cerebrospinal fluid in Alzheimer’s disease. Nature. 2020;577(7790):399–404.

40. Lepennetier G, Hracsko Z, Unger M, Van Griensven M, Grummel V, Krumbholz M, et al. Cytokine and immune cell profiling in the cerebrospinal fluid of patients with neuro-inflammatory diseases. J Neuroinflammation. 2019;16(1):219.

41. Plog BA, Dashnaw ML, Hitomi E, Peng W, Liao Y, Lou N, et al. Biomarkers of traumatic injury are transported from brain to blood via the glymphatic system. J Neurosci. 2015;35(2):518–26.

42. Ringstad G, Eide PK. Cerebrospinal fluid tracer efflux to parasagittal dura in humans. Nat Commun. 2020;11(1):354.

43. Louveau A, Plog BA, Antila S, Alitalo K, Nedergaard M, Kipnis J. Understanding the functions and relationships of the glymphatic system and meningeal lymphatics. J Clin Invest. 2017;127(9):3210–9.

44. Goodman JR, Iliff JJ. Vasomotor influences on glymphatic-lymphatic coupling and solute trafficking in the central nervous system. J Cereb Blood Flow Metab. 2019:271678X19874134.

45. Antila S, Karaman S, Nurmi H, Airavaara M, Voutilainen MH, Mathivet T, et al. Development and plasticity of meningeal lymphatic vessels. J Exp Med. 2017;214(12):3645–67.

46. Iliff JJ, Wang M, Zeppenfeld DM, Venkataraman A, Plog BA, Liao Y, et al. Cerebral arterial pulsation drives paravascular CSF-interstitial fluid exchange in the murine brain. J Neurosci. 2013;33(46):18190–9.

47. Iliff JJ, Chen MJ, Plog BA, Zeppenfeld DM, Soltero M, Yang L, et al. Impairment of glymphatic pathway function promotes tau pathology after traumatic brain injury. J Neurosci. 2014;34(49):16180–93.

48. Iliff JJ, Wang M, Liao Y, Plogg BA, Peng W, Gundersen GA, et al. A paravascular pathway facilitates CSF flow through the brain parenchyma and the clearance of interstitial solutes, including amyloid beta. Sci Transl Med. 2012;4(147):147ra11.

49. Jessen NA, Munk AS, Lundgaard I, Nedergaard M. The Glymphatic System: A Beginner’s Guide. Neurochem Res. 2015;40(12):2583–99.

50. Peng W, Achariyar TM, Li B, Liao Y, Mestre H, Hitomi E, et al. Suppression of glymphatic fluid transport in a mouse model of Alzheimer’s disease. Neurobiol Dis. 2016;93:215–25.

51. Ren Z, Iliff JJ, Yang L, Yang J, Chen X, Chen MJ, et al. ’Hit & Run’ model of closed-skull traumatic brain injury (TBI) reveals complex patterns of post-traumatic AQP4 dysregulation. J Cereb Blood Flow Metab. 2013;33(6):834–45.

52. Bolte AC, Dutta AB, Hurt ME, Smirnov I, Kovacs MA, McKee CA, et al. Meningeal lymphatic dysfunction exacerbates traumatic brain injury pathogenesis. Nat Commun. 2020;11(1):4524.

53. Stuart T, Butler A, Hoffman P, Hafemeister C, Papalexi E, Mauck WM, 3rd, et al. Comprehensive Integration of Single-Cell Data. Cell. 2019;177(7):1888–902 e21.

54. Butler A, Hoffman P, Smibert P, Papalexi E, Satija R. Integrating single-cell transcriptomic data across different conditions, technologies, and species. Nat Biotechnol. 2018;36(5):411–20.

55. Shao X, Liao J, Lu X, Xue R, Ai N, Fan X. scCATCH: Automatic Annotation on Cell Types of Clusters from Single-Cell RNA Sequencing Data. iScience. 2020;23(3):100882.

56. Risso D, Perraudeau F, Gribkova S, Dudoit S, Vert JP. A general and flexible method for signal extraction from single-cell RNA-seq data. Nat Commun. 2018;9(1):284.

57. Shannon P, Markiel A, Ozier O, Baliga NS, Wang JT, Ramage D, et al. Cytoscape: a software environment for integrated models of biomolecular interaction networks. Genome Res. 2003;13(11):2498–504.

58. Chen J, Xu H, Aronow BJ, Jegga AG. Improved human disease candidate gene prioritization using mouse phenotype. BMC Bioinformatics. 2007;8:392.

59. Wickham H. ggplot2: Elegant Graphics for Data Analysis: Springer-Verlag New York; 2016.

60. Wickham H, Averick M, Bryan J, Chang W, McGowan L, François R, et al. Welcome to the Tidyverse. Journal of Open Source Software. 2019;4(43).

61. Trapnell C, Cacchiarelli D, Grimsby J, Pokharel P, Li S, Morse M, et al. The dynamics and regulators of cell fate decisions are revealed by pseudotemporal ordering of single cells. Nat Biotechnol. 2014;32(4):381–6.

62. Kim D, Paggi JM, Park C, Bennett C, Salzberg SL. Graph-based genome alignment and genotyping with HISAT2 and HISAT-genotype. Nat Biotechnol. 2019;37(8):907–15.

63. Li H, Handsaker B, Wysoker A, Fennell T, Ruan J, Homer N, et al. The Sequence Alignment/Map format and SAMtools. Bioinformatics. 2009;25(16):2078–9.

64. Anders S, Pyl PT, Huber W. HTSeq--a Python framework to work with high-throughput sequencing data. Bioinformatics. 2015;31(2):166–9.

65. Love MI, Huber W, Anders S. Moderated estimation of fold change and dispersion for RNA-seq data with DESeq2. Genome Biol. 2014;15(12):550.

66. Liberzon A, Birger C, Thorvaldsdottir H, Ghandi M, Mesirov JP, Tamayo P. The Molecular Signatures Database (MSigDB) hallmark gene set collection. Cell Syst. 2015;1(6):417–25.

67. Haimon Z, Volaski A, Orthgiess J, Boura-Halfon S, Varol D, Shemer A, et al. Re-evaluating microglia expression profiles using RiboTag and cell isolation strategies. Nat Immunol. 2018;19(6):636–44.

68. Marsh SE, Kamath T, Walker AJ, Dissing-Olesen L, Hammond TR, Young AMH, et al. Single cell sequencing reveals glial specific responses to tissue processing and enzymatic dissocation in mice and humans. 2020.

69. Van Hove H, Martens L, Scheyltjens I, De Vlaminck K, Pombo Antunes AR, De Prijck S, et al. A single-cell atlas of mouse brain macrophages reveals unique transcriptional identities shaped by ontogeny and tissue environment. Nat Neurosci. 2019;22(6):1021–35.

70. Karve IP, Zhang M, Habgood M, Frugier T, Brody KM, Sashindranath M, et al. Ablation of Type-1 IFN Signaling in Hematopoietic Cells Confers Protection Following Traumatic Brain Injury. eNeuro. 2016;3(1).

71. Barrett JP, Henry RJ, Shirey KA, Doran SJ, Makarevich OD, Ritzel RM, et al. Interferon-beta Plays a Detrimental Role in Experimental Traumatic Brain Injury by Enhancing Neuroinflammation That Drives Chronic Neurodegeneration. J Neurosci. 2020;40(11):2357–70.

72. Park SY, Jung MY, Lee SJ, Kang KB, Gratchev A, Riabov V, et al. Stabilin-1 mediates phosphatidylserine-dependent clearance of cell corpses in alternatively activated macrophages. J Cell Sci. 2009;122(Pt 18):3365–73.

73. Noubade R, Wong K, Ota N, Rutz S, Eidenschenk C, Valdez PA, et al. NRROS negatively regulates reactive oxygen species during host defence and autoimmunity. Nature. 2014;509(7499):235–9.

74. Hung WS, Ling P, Cheng JC, Chang SS, Tseng CP. Disabled-2 is a negative immune regulator of lipopolysaccharide-stimulated Toll-like receptor 4 internalization and signaling. Sci Rep. 2016;6:35343.

75. Stables MJ, Shah S, Camon EB, Lovering RC, Newson J, Bystrom J, et al. Transcriptomic analyses of murine resolution-phase macrophages. Blood. 2011;118(26):e192–208.

76. Meyaard L, Adema GJ, Chang C, Woollatt E, Sutherland GR, Lanier LL, et al. LAIR-1, a Novel Inhibitory Receptor Expressed on Human Mononuclear Leukocytes. Immunity. 1997;7(2):283–90.

77. Kwiecien I, Polubiec-Kownacka M, Dziedzic D, Wolosz D, Rzepecki P, Domagala-Kulawik J. CD163 and CCR7 as markers for macrophage polarization in lung cancer microenvironment. Cent Eur J Immunol. 2019;44(4):395–402.

78. Martinez FO, Gordon S. The M1 and M2 paradigm of macrophage activation: time for reassessment. F1000Prime Rep. 2014;6:13.

79. DeSisto J, O’Rourke R, Jones HE, Pawlikowski B, Malek AD, Bonney S, et al. Single-Cell Transcriptomic Analyses of the Developing Meninges Reveal Meningeal Fibroblast Diversity and Function. Dev Cell. 2020;54(1):43–59 e4.

80. Doro D, Liu A, Grigoriadis AE, Liu KJ. The Osteogenic Potential of the Neural Crest Lineage May Contribute to Craniosynostosis. Mol Syndromol. 2019;10(1-2):48–57.

81. Cooper GM, Durham EL, Cray JJ, Jr., Siegel MI, Losee JE, Mooney MP. Tissue interactions between craniosynostotic dura mater and bone. J Craniofac Surg. 2012;23(3):919–24.

82. Zarbalis K, Siegenthaler JA, Choe Y, May SR, Peterson AS, Pleasure SJ. Cortical dysplasia and skull defects in mice with a Foxc1 allele reveal the role of meningeal differentiation in regulating cortical development. Proc Natl Acad Sci U S A. 2007;104(35):14002–7.

83. Kalamarides M, Stemmer-Rachamimov AO, Niwa-Kawakita M, Chareyre F, Taranchon E, Han ZY, et al. Identification of a progenitor cell of origin capable of generating diverse meningioma histological subtypes. Oncogene. 2011;30(20):2333–44.

84. Siegenthaler JA, Ashique AM, Zarbalis K, Patterson KP, Hecht JH, Kane MA, et al. Retinoic acid from the meninges regulates cortical neuron generation. Cell. 2009;139(3):597–609.

85. Caglayan AO, Baranoski JF, Aktar F, Han W, Tuysuz B, Guzel A, et al. Brain malformations associated with Knobloch syndrome--review of literature, expanding clinical spectrum, and identification of novel mutations. Pediatr Neurol. 2014;51(6):806–13 e8.

86. Rua R, McGavern DB. Advances in Meningeal Immunity. Trends Mol Med. 2018;24(6):542–59.

87. Tibbitt CA, Stark JM, Martens L, Ma J, Mold JE, Deswarte K, et al. Single-Cell RNA Sequencing of the T Helper Cell Response to House Dust Mites Defines a Distinct Gene Expression Signature in Airway Th2 Cells. Immunity. 2019;51(1):169–84 e5.

88. Hu D, Notarbartolo S, Croonenborghs T, Patel B, Cialic R, Yang TH, et al. Transcriptional signature of human pro-inflammatory TH17 cells identifies reduced IL10 gene expression in multiple sclerosis. Nat Commun. 2017;8(1):1600.

89. de Winde CM, Veenbergen S, Young KH, Xu-Monette ZY, Wang XX, Xia Y, et al. Tetraspanin CD37 protects against the development of B cell lymphoma. J Clin Invest. 2016;126(2):653–66.

90. Lee RD, Munro SA, Knutson TP, LaRue RS, Heltemes-Harris LM, Farrar MA. Single-cell analysis of developing B cells reveals dynamic gene expression networks that govern B cell development and transformation. 2020.

91. Liau NPD, Laktyushin A, Lucet IS, Murphy JM, Yao S, Whitlock E, et al. The molecular basis of JAK/STAT inhibition by SOCS1. Nat Commun. 2018;9(1):1558.

92. Toh BH, Kyaw T, Tipping P, Bobik A. Immune regulation by CD52-expressing CD4 T cells. Cell Mol Immunol. 2013;10(5):379–82.

93. Fitzpatrick Z, Frazer G, Ferro A, Clare S, Bouladoux N, Ferdinand J, et al. Gut-educated IgA plasma cells defend the meningeal venous sinuses. Nature. 2020;587(7834):472–6.

94. Brioschi S, Wang WL, Peng V, Wang M, Shchukina I, Greenberg ZJ, et al. Heterogeneity of meningeal B cells reveals a lymphopoietic niche at the CNS borders. Science. 2021;373(6553).

95. Mrdjen D, Pavlovic A, Hartmann FJ, Schreiner B, Utz SG, Leung BP, et al. High-Dimensional Single-Cell Mapping of Central Nervous System Immune Cells Reveals Distinct Myeloid Subsets in Health, Aging, and Disease. Immunity. 2018;48(2):380–95 e6.

96. Baruch K, Deczkowska A, David E, Castellano JM, Miller O, Kertser A, et al. Aging. Aging-induced type I interferon response at the choroid plexus negatively affects brain function. Science. 2014;346(6205):89–93.

97. Abdullah A, Zhang M, Frugier T, Bedoui S, Taylor JM, Crack PJ. STING-mediated type-I interferons contribute to the neuroinflammatory process and detrimental effects following traumatic brain injury. J Neuroinflammation. 2018;15(1):323.

98. Zhang M, Downes CE, Wong CHY, Brody KM, Guio-Agulair PL, Gould J, et al. Type-I interferon signalling through IFNAR1 plays a deleterious role in the outcome after stroke. Neurochem Int. 2017;108:472–80.

99. Louveau A, Herz J, Alme MN, Salvador AF, Dong MQ, Viar KE, et al. CNS lymphatic drainage and neuroinflammation are regulated by meningeal lymphatic vasculature. Nature Neuroscience. 2018;21(10):1380–91.

100. Yang AC, Stevens MY, Chen MB, Lee DP, Stahli D, Gate D, et al. Physiological blood-brain transport is impaired with age by a shift in transcytosis. Nature. 2020;583(7816):425–30.

101. Fitzgerald-Bocarsly P, Dai J, Singh S. Plasmacytoid dendritic cells and type I IFN: 50 years of convergent history. Cytokine Growth Factor Rev. 2008;19(1):3–19.

102. Gupte R, Brooks W, Vukas R, Pierce J, Harris J. Sex Differences in Traumatic Brain Injury: What We Know and What We Should Know. J Neurotrauma. 2019;36(22):3063–91.

103. Marschallinger J, Iram T, Zardeneta M, Lee SE, Lehallier B, Haney MS, et al. Lipid-droplet-accumulating microglia represent a dysfunctional and proinflammatory state in the aging brain. Nat Neurosci. 2020;23(2):194–208.

104. Itoh T, Imano M, Nishida S, Tsubaki M, Mizuguchi N, Hashimoto S, et al. Increased apoptotic neuronal cell death and cognitive impairment at early phase after traumatic brain injury in aged rats. Brain Struct Funct. 2013;218(1):209–20.

105. Chen J, Wang L, Xu H, Xing L, Zhuang Z, Zheng Y, et al. Meningeal lymphatics clear erythrocytes that arise from subarachnoid hemorrhage. Nat Commun. 2020;11(1):3159.

106. Pu T, Zou W, Feng W, Zhang Y, Wang L, Wang H, et al. Persistent Malfunction of Glymphatic and Meningeal Lymphatic Drainage in a Mouse Model of Subarachnoid Hemorrhage. Exp Neurobiol. 2019;28(1):104–18.

107. Da Mesquita S, Louveau A, Vaccari A, Smirnov I, Cornelison RC, Kingsmore KM, et al. Functional aspects of meningeal lymphatics in ageing and Alzheimer’s disease. Nature. 2018;560(7717):185–91.

108. Ahn JH, Cho H, Kim JH, Kim SH, Ham JS, Park I, et al. Meningeal lymphatic vessels at the skull base drain cerebrospinal fluid. Nature. 2019.

109. Ma Q, Ineichen BV, Detmar M, Proulx ST. Outflow of cerebrospinal fluid is predominantly through lymphatic vessels and is reduced in aged mice. Nat Commun. 2017;8(1):1434.

