## Supplementary material for "The meningeal transcriptional response to traumatic brain injury and aging": Supp Figures

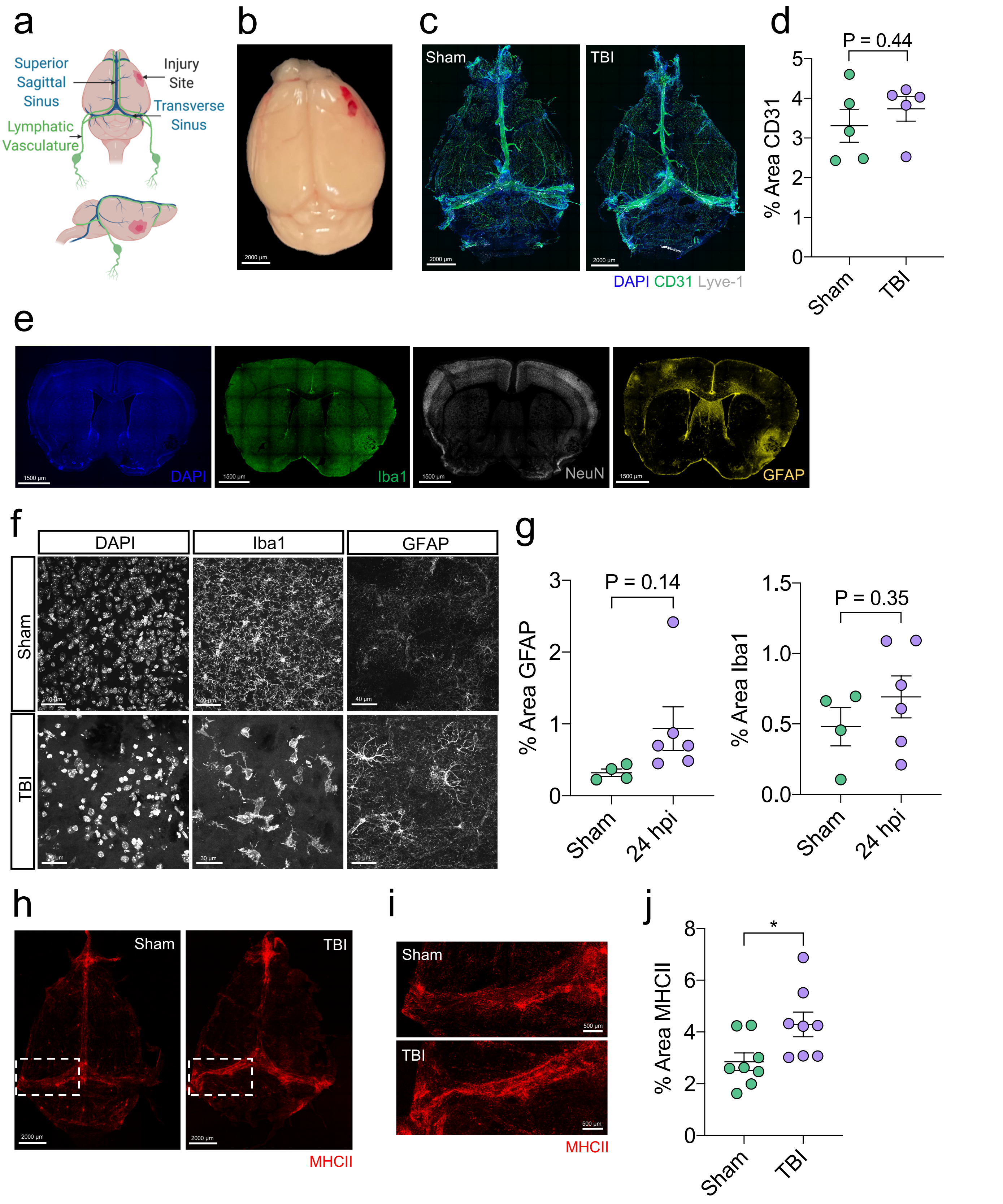


Supplementary Figure 1

a

### Activated Macs 1

| Gene | P_adj |
|---|---|
| C1qb | 0 |
| C1qc | 0 |
| C1qa | 0 |
| Lyz2 | 0 |
| Csf1r | 0 |
| Ctss | 0 |
| Ctsc | 0 |
| Lgmn | 0 |
| Ms4a7 | 0 |
| Hexb | 0 |
| Mrc1 | 0 |
| Cx3cr1 | 0 |
| Fcrls | 0 |
| Fcgr3 | 0 |
| Mgl2 | 0 |
| Cd68 | 0 |
| Pf4 | 0 |
| C3ar1 | 0 |
| Fcer1g | 0 |
| Adgre1 | 0 |

### Activated Macs 2

| Gene | P_adj |
|---|---|
| C1qa | 1.69E-267 |
| Pf4 | 3.75E-257 |
| Cbr2 | 4.59E-253 |
| C1qc | 1.35E-251 |
| C1qb | 5.57E-237 |
| F13a1 | 1.38E-195 |
| Lyz2 | 3.18E-177 |
| Apoe | 8.48E-157 |
| Ccl8 | 1.16E-148 |
| Mrc1 | 5.64E-131 |
| Fcer1g | 3.84E-130 |
| Ms4a7 | 5.96E-130 |
| Csf1r | 2.62E-126 |
| Fcrls | 2.45E-122 |
| Ftl1 | 3.90E-122 |
| Selenop | 2.68E-108 |
| Tyrobp | 4.08E-108 |
| Stab1 | 2.42E-94 |
| Folr2 | 9.74E-92 |
| Hpgd | 2.65E-89 |

### Macs 3

| Gene | P_adj |
|---|---|
| Apol7c | 0 |
| Cacnb3 | 0 |
| Ccl22 | 0 |
| Nudt17 | 0 |
| Gm13546 | 0 |
| Gm10851 | 0 |
| Slco5a1 | 0 |
| Mreg | 0 |
| Lad1 | 0 |
| Il12b | 3.12E-296 |
| H2-M2 | 2.19E-293 |
| Gm47662 | 1.42E-274 |
| Arc | 2.89E-257 |
| Tmem150c | 4.64E-233 |
| Mmp25 | 4.31E-205 |
| Cacna1s | 4.46E-204 |
| Zmynd15 | 1.82E-187 |
| Bcl2l14 | 3.06E-177 |
| Il4i1 | 5.84E-169 |
| Cdkn2b | 4.73E-168 |

### B Cells 1

| Gene | P_adj |
|---|---|
| Ms4a1 | 0 |
| Ly6d | 0 |
| Cd79a | 0 |
| Cd79b | 0 |
| Igkc | 0 |
| Iglc2 | 0 |
| Iglc1 | 0 |
| Vpreb3 | 0 |
| Tnfrsf13c | 0 |
| Iglc3 | 0 |
| Cd24a | 0 |
| Spib | 0 |
| Siglecg | 0 |
| Fcmr | 0 |
| Cd19 | 0 |
| Fcrla | 0 |
| Fam129c | 0 |
| Pax5 | 0 |
| Pou2af1 | 0 |
| Srpk3 | 0 |

### B Cells 2

| Gene | P_adj |
|---|---|
| Cd79a | 5.12E-295 |
| Iglc2 | 5.77E-292 |
| Ighd | 3.91E-282 |
| Fcmr | 5.28E-281 |
| Fcer2a | 2.85E-276 |
| Ms4a1 | 1.87E-260 |
| Cd79b | 1.85E-221 |
| Ly6d | 9.85E-221 |
| Iglc3 | 1.44E-205 |
| Igkc | 5.27E-191 |
| Cd55 | 4.09E-174 |
| Cd19 | 9.60E-166 |
| H2-DMb2 | 3.26E-159 |
| H2-Ob | 1.35E-148 |
| Cd37 | 1.71E-144 |
| Bank1 | 8.25E-139 |
| Ighm | 5.34E-123 |
| Cxcr5 | 4.08E-122 |
| Siglecg | 4.68E-119 |
| Ebf1 | 7.44E-119 |

### Immature/Diff B Cells

| Gene | P_adj |
|---|---|
| Rag1 | 0 |
| Cecr2 | 0 |
| Rag2 | 0 |
| Cplx2 | 1.72E-298 |
| AL611986.1 | 2.83E-293 |
| Gm37065 | 4.82E-277 |
| Vpreb3 | 1.77E-269 |
| Myb | 3.91E-267 |
| Slc12a3 | 1.55E-254 |
| Xrcc6 | 1.53E-233 |
| Arl5c | 1.09E-207 |
| Pou2af1 | 2.05E-206 |
| Myl4 | 2.00E-204 |
| Fam129c | 1.39E-199 |
| Uchl1 | 9.52E-191 |
| Bach2 | 2.52E-189 |
| Fcrla | 8.19E-188 |
| Bcl7a | 2.76E-177 |
| Spns3 | 4.15E-170 |
| Pafah1b3 | 1.07E-168 |

### CD3+ T Cells

| Gene | P_adj |
|---|---|
| Cd3g | 0 |
| Cd3e | 0 |
| Cd3d | 0 |
| Ms4a4b | 0 |
| Thy1 | 0 |
| Cxcr6 | 0 |
| Nkg7 | 0 |
| Ctsw | 0 |
| Xcl1 | 0 |
| Hcst | 0 |
| Gimap3 | 0 |
| Lck | 0 |
| Il2rb | 0 |
| Gimap4 | 0 |
| Lat | 0 |
| Cxcr3 | 0 |
| Skap1 | 0 |
| Trbc2 | 0 |
| Cd247 | 0 |
| Trac | 0 |

### Activated T Cells

| Gene | P_adj |
|---|---|
| Gata3 | 0 |
| Rnf128 | 0 |
| Il7r | 1.80E-278 |
| Il1rl1 | 5.87E-247 |
| Skap1 | 2.61E-216 |
| Cxcr6 | 3.34E-201 |
| Inpp4b | 4.90E-196 |
| Tnfsf11 | 2.89E-158 |
| Thy1 | 8.90E-158 |
| Faah | 1.25E-153 |
| Tespa1 | 8.07E-145 |
| Ltb4r1 | 5.18E-134 |
| Il2ra | 4.71E-132 |
| Lat | 8.00E-130 |
| Gimap3 | 1.24E-117 |
| Bcl11b | 2.06E-116 |
| Cish | 1.79E-115 |
| Ltb | 8.29E-108 |
| Icos | 1.74E-107 |
| Itk | 1.94E-100 |

### NK Cells

| Gene | P_adj |
|---|---|
| Gzma | 0 |
| Nkg7 | 0 |
| Gzmb | 0 |
| Ncr1 | 0 |
| Prf1 | 0 |
| Klrb1c | 0 |
| Klre1 | 0 |
| Klra8 | 0 |
| Klra9 | 0 |
| Klra4 | 0 |
| Klra7 | 0 |
| Klrc2 | 0 |
| Eomes | 0 |
| Klri2 | 0 |
| S1pr5 | 0 |
| Gm19585 | 6.12E-298 |
| Ctsw | 1.57E-277 |
| Adamts14 | 5.46E-277 |
| Il2rb | 2.67E-262 |
| Ccl5 | 1.28E-258 |

### Dendritic Cells

| Gene | P_adj |
|---|---|
| Cd209a | 0 |
| Plbd1 | 0 |
| Cd209c | 0 |
| Olfm1 | 0 |
| Jaml | 0 |
| Flt3 | 0 |
| Mcemp1 | 1.08E-299 |
| Skint3 | 1.81E-298 |
| Lgals3 | 2.54E-280 |
| Ear2 | 1.40E-271 |
| Slamf7 | 5.67E-263 |
| Dna2 | 7.81E-245 |
| Lmo1 | 1.91E-242 |
| Hepacam2 | 3.34E-236 |
| Itgax | 7.57E-235 |
| H2-DMb1 | 1.56E-232 |
| Rtl5 | 1.05E-229 |
| Ltb4r1 | 7.19E-229 |
| Ciita | 4.96E-228 |
| Slamf8 | 4.69E-225 |

### Plasmacytoid Dendritic Cells

| Gene | P_adj |
|---|---|
| Cox6a2 | 0 |
| Siglech | 0 |
| Klk1 | 0 |
| Ccr9 | 0 |
| AC140186.1 | 0 |
| Gm21762 | 0 |
| Cd300c | 0 |
| Klra17 | 0 |
| Atp2a1 | 0 |
| Dntt | 0 |
| Foxr1 | 0 |
| Sh3bgr | 0 |
| Grm8 | 0 |
| Gm12253 | 0 |
| Havcr1 | 0 |
| Pacsin1 | 7.60E-293 |
| Lefty1 | 1.08E-292 |
| Ccdc162 | 6.99E-291 |
| Pdzd4 | 7.40E-281 |
| Cd209d | 5.59E-264 |

### Neutrophils

| Gene | P_adj |
|---|---|
| Retnlg | 0 |
| Lcn2 | 0 |
| Slpi | 0 |
| Hp | 0 |
| Mmp8 | 0 |
| Cxcr2 | 0 |
| Ly6g | 0 |
| Hdc | 0 |
| Hcar2 | 0 |
| Padi4 | 0 |
| Trem3 | 0 |
| Cebpe | 0 |
| Itgb2l | 0 |
| Ankrd22 | 0 |
| Cd177 | 2.35E-259 |
| Chil1 | 9.41E-259 |
| Wfdc21 | 4.83E-238 |
| Mmp9 | 7.38E-232 |
| Trem1 | 5.82E-223 |
| Slfn4 | 2.79E-220 |

Supplementary Figure 2a

b

### Proliferating Cells

| Gene | P_adj |
|---|---|
| Mki67 | 0 |
| Top2a | 0 |
| Pclaf | 0 |
| Cenpf | 0 |
| Birc5 | 0 |
| Rrm2 | 0 |
| Ube2c | 0 |
| Ccnb2 | 0 |
| Cdca8 | 0 |
| Ccna2 | 0 |
| Tpx2 | 0 |
| Cenpe | 0 |
| Nusap1 | 0 |
| Knl1 | 0 |
| Cdca3 | 0 |
| Aspm | 0 |
| Kif11 | 0 |
| Cdk1 | 0 |
| Pbk | 0 |
| Spc24 | 0 |

### Fibroblasts

| Gene | P_adj |
|---|---|
| Col1a1 | 0 |
| Igfbp5 | 0 |
| Col1a2 | 0 |
| Dcn | 0 |
| Slc38a2 | 0 |
| Col12a1 | 0 |
| Mfap4 | 0 |
| Slc47a1 | 0 |
| Serpinf1 | 0 |
| Snhg11 | 0 |
| Slc4a10 | 0 |
| Sfrp4 | 0 |
| Mgp | 0 |
| Pcolce | 0 |
| Fbln1 | 0 |
| Cdh11 | 0 |
| Col6a2 | 0 |
| Slc23a2 | 0 |
| Igfbp6 | 0 |
| Ank2 | 0 |

### Endothelial Cells 1

| Gene | P_adj |
|---|---|
| Igfbp3 | 0 |
| Ly6c1 | 0 |
| Plvap | 0 |
| Flt1 | 0 |
| Kdr | 0 |
| Plpp1 | 0 |
| Emcn | 0 |
| Rgcc | 0 |
| Podxl | 0 |
| Ptprb | 0 |
| Gpihbp1 | 0 |
| Egfl7 | 0 |
| Igfbp7 | 0 |
| Ly6a | 0 |
| Slc9a3r2 | 0 |
| Sparcl1 | 0 |
| Scarb1 | 0 |
| Tm4sf1 | 0 |
| Pecam1 | 0 |
| Hspb1 | 0 |

### Endothelial Cells 2

| Gene | P_adj |
|---|---|
| Vwf | 0 |
| Lrg1 | 0 |
| Ptprb | 3.41E-284 |
| Clec14a | 2.41E-282 |
| Slco2a1 | 6.38E-267 |
| Wipf3 | 1.57E-255 |
| Tgm2 | 1.67E-254 |
| Flt1 | 1.28E-229 |
| Egfl7 | 2.01E-229 |
| Adgrg6 | 1.10E-221 |
| Nos3 | 4.73E-216 |
| Nrp2 | 8.75E-196 |
| Cavin2 | 1.74E-190 |
| Cldn5 | 6.70E-184 |
| Fam174b | 2.31E-178 |
| Cfh | 8.64E-178 |
| Mmrn2 | 4.39E-169 |
| Gpr182 | 5.45E-168 |
| AU021092 | 1.38E-167 |
| Tek | 2.35E-166 |

### Pericytes

| Gene | P_adj |
|---|---|
| Rgs5 | 0 |
| Notch3 | 0 |
| Abcc9 | 0 |
| Myh11 | 0 |
| Ndufa4l2 | 0 |
| Higd1b | 0 |
| Rgs4 | 0 |
| Gucy1b1 | 2.57E-289 |
| Trpc3 | 2.11E-277 |
| Itga7 | 3.32E-266 |
| Steap4 | 1.62E-262 |
| Rasl12 | 2.47E-258 |
| Ano1 | 1.12E-237 |
| Kcnj8 | 1.12E-227 |
| Foxs1 | 3.69E-216 |
| Cox4i2 | 2.84E-214 |
| Rasl11a | 2.19E-174 |
| Aspn | 1.11E-156 |
| Pdgfrb | 1.63E-156 |
| Rem1 | 7.97E-155 |

### Choroid Plexus

| Gene | P_adj |
|---|---|
| Enpp2 | 0 |
| Mt3 | 0 |
| Ldhb | 0 |
| 2900040C04Rik | 0 |
| Ppp1r1b | 0 |
| Calml4 | 0 |
| Kl | 0 |
| Sostdc1 | 0 |
| Car2 | 0 |
| Clic6 | 0 |
| Prr32 | 0 |
| Gsta4 | 0 |
| Krt18 | 0 |
| Hemk1 | 0 |
| Folr1 | 0 |
| Kcnj13 | 0 |
| Fam81a | 0 |
| F5 | 0 |
| Kcne2 | 0 |
| Drc7 | 0 |

### Pineal Gland Cells

| Gene | P_adj |
|---|---|
| Pde6g | 0 |
| Gnb3 | 0 |
| Mgarp | 0 |
| Snap25 | 0 |
| Bsx | 0 |
| Pax6 | 1.19E-298 |
| Spink4 | 7.29E-285 |
| Crabp1 | 1.50E-282 |
| Tph1 | 2.56E-271 |
| Cplx3 | 8.17E-271 |
| Chgb | 9.10E-262 |
| Cdhr1 | 8.06E-252 |
| Pcsk1n | 1.75E-225 |
| Bcat1 | 1.80E-220 |
| Cspg5 | 6.41E-210 |
| Plekhb1 | 2.34E-205 |
| Syp | 9.34E-168 |
| Tst | 1.48E-159 |
| Cntnap2 | 1.16E-158 |
| Edn3 | 1.95E-149 |

### Stem Cells

| Gene | P_adj |
|---|---|
| Mpz | 0 |
| Kcna1 | 0 |
| Plp1 | 0 |
| Scn7a | 0 |
| Mal | 0 |
| Gpm6b | 0 |
| Gfra3 | 0 |
| Kcna2 | 0 |
| Aatk | 0 |
| Cmtm5 | 0 |
| Cdh19 | 0 |
| Art3 | 0 |
| Egfl8 | 0 |
| Gpr37l1 | 0 |
| Abca8a | 0 |
| Lgi4 | 0 |
| Sox10 | 0 |
| Ank3 | 0 |
| Chl1 | 0 |
| Abca8b | 0 |

### Clotting Related

| Gene | P_adj |
|---|---|
| F10 | 5.87E-299 |
| Treml4 | 2.64E-247 |
| Hp | 7.00E-211 |
| Gm9733 | 4.60E-198 |
| Sirpb1c | 1.28E-197 |
| Ifitm6 | 2.73E-183 |
| Nfe2 | 7.37E-176 |
| Adgre4 | 2.09E-163 |
| Emilin2 | 3.60E-143 |
| Atp1a3 | 1.42E-116 |
| Fgr | 7.51E-112 |
| Gsr | 6.44E-111 |
| Ear2 | 1.71E-98 |
| Cebpb | 1.05E-79 |
| Gda | 1.31E-74 |
| Sirpb1b | 2.54E-72 |
| Slc16a3 | 1.37E-71 |
| Plac8 | 7.90E-71 |
| Ace | 2.63E-70 |
| Tnfrsf1b | 2.66E-69 |

Supplementary Figure 2b

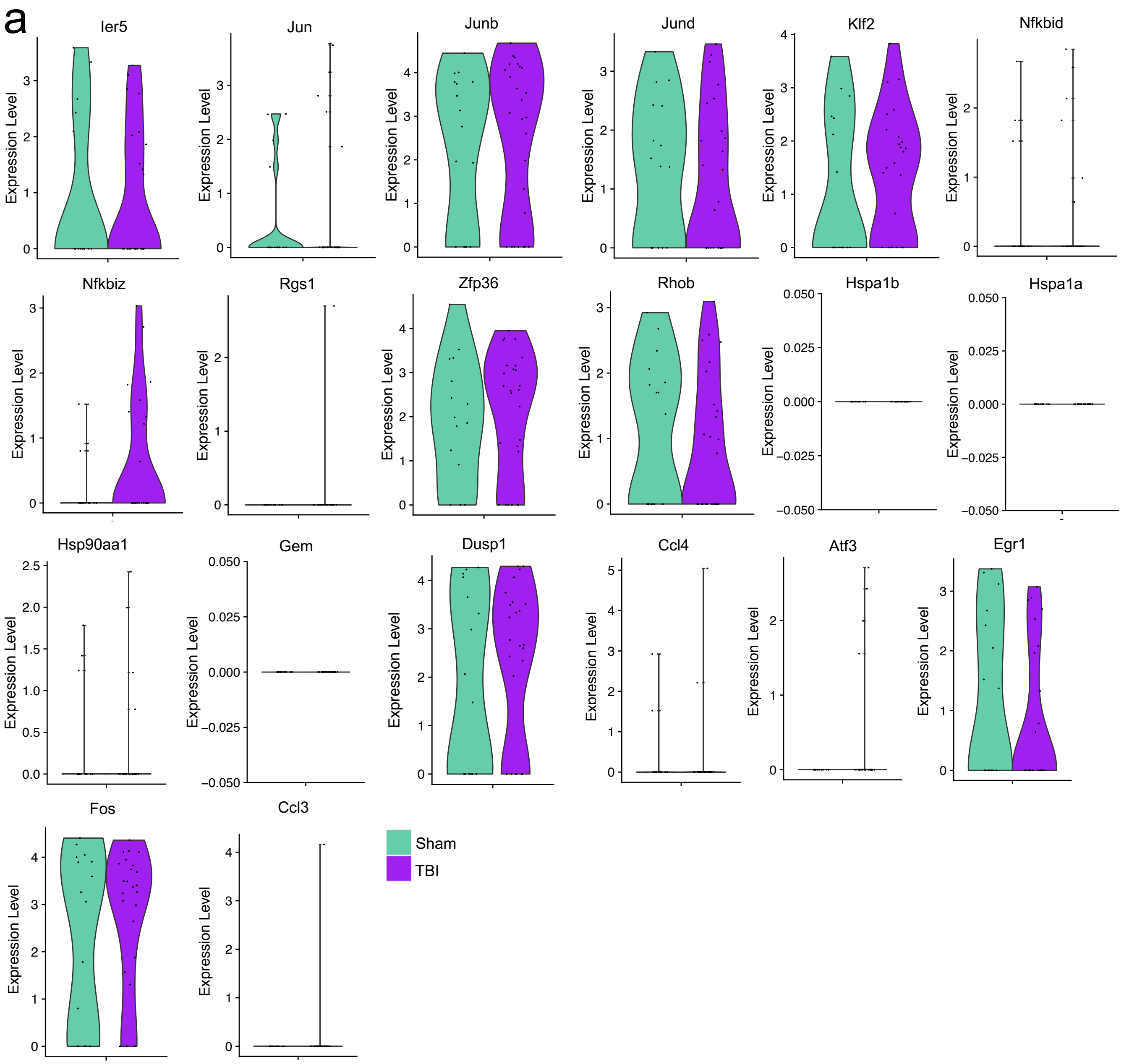


Supplementary Figure 3

a

Ferritin Expressing

| Gene | P_adj |
|---|---|
| Ftl1 | 2.28E-88 |
| Fth1 | 3.91E-53 |
| Pf4 | 4.42E-38 |
| Rps29 | 8.98E-35 |
| Ccl8 | 3.67E-34 |
| Selenop | 5.36E-34 |
| Rplp1 | 3.02E-32 |
| Cbr2 | 2.54E-29 |
| Tyrobp | 1.77E-24 |
| Igfbp4 | 1.27E-22 |

Anti-Inflammatory

| Gene | P_adj |
|---|---|
| Mrc1 | 3.93E-58 |
| C5ar1 | 9.21E-56 |
| Stab1 | 3.55E-51 |
| Nrros | 3.14E-46 |
| Pmp22 | 9.12E-45 |
| Slc9a9 | 6.64E-44 |
| Fcrls | 1.59E-42 |
| Maf | 2.14E-42 |
| Dab2 | 3.78E-42 |
| Rab11fip5 | 1.18E-41 |

Resolution Phase

| Gene | P_adj |
|---|---|
| F11r | 3.81E-71 |
| Olfml3 | 1.73E-67 |
| Ctss | 1.81E-66 |
| Hexb | 8.61E-59 |
| Tgfbr1 | 8.97E-56 |
| Slamf9 | 7.60E-48 |
| Psap | 2.32E-44 |
| Mafb | 1.25E-41 |
| H2-Eb1 | 1.35E-41 |
| Lair1 | 4.44E-41 |

Inflammatory 1

| Gene | P_adj |
|---|---|
| Napsa | 2.45E-60 |
| S100a6 | 4.68E-57 |
| Capg | 4.86E-46 |
| S100a4 | 2.96E-45 |
| Lgals3 | 8.75E-39 |
| Fgr | 9.46E-39 |
| Vim | 2.93E-34 |
| Tnip3 | 3.09E-33 |
| Ccr2 | 2.56E-32 |
| Alcam | 2.63E-30 |

Inflammatory 2

| Gene | P_adj |
|---|---|
| Ccr7 | 2.23E-235 |
| Cacnb3 | 7.88E-201 |
| Nudt17 | 2.44E-164 |
| Socs2 | 2.16E-159 |
| Spint2 | 7.69E-148 |
| Mreg | 2.60E-137 |
| Slco5a1 | 1.21E-128 |
| Gm10851 | 8.54E-120 |
| Apol7c | 8.22E-118 |
| Serpinb6b | 4.47E-113 |

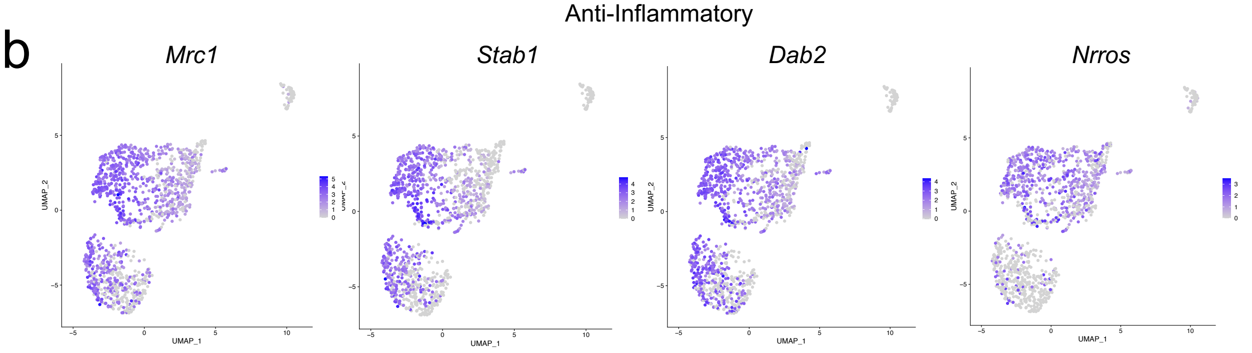


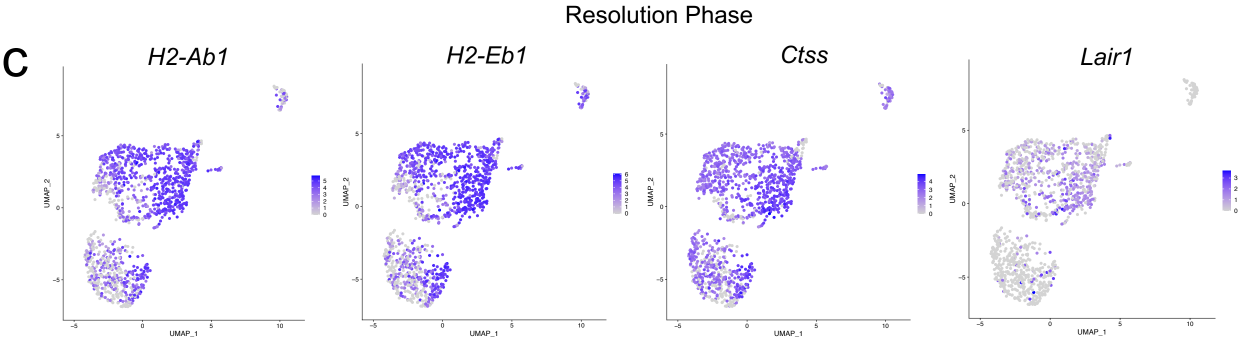


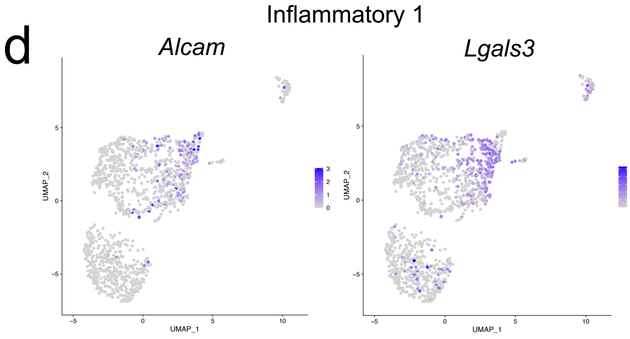


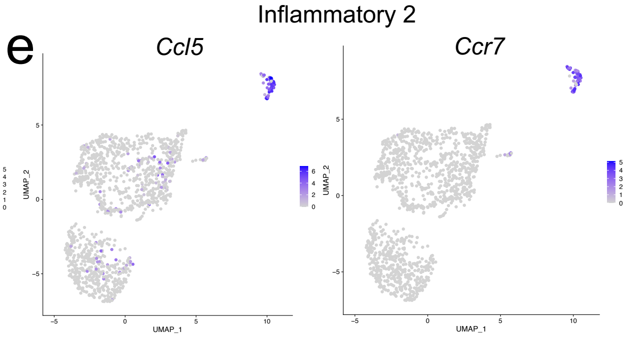


Supplementary Figure 4

a

CD8+ T Cells

| Gene | P_adj |
|---|---|
| Ms4a6b | 7.05E-44 |
| Ifi27l2a | 8.59E-44 |
| Ms4a4b | 3.49E-43 |
| Lck | 1.15E-41 |
| Slfn1 | 7.64E-39 |
| Cd3e | 4.95E-37 |
| Cd3g | 7.77E-36 |
| Cd2 | 1.88E-32 |
| Cd3d | 1.52E-31 |
| Cd28 | 2.75E-31 |

Th2 Cells

| Gene | P_adj |
|---|---|
| Lpcat2 | 1.37E-68 |
| Rnf128 | 1.59E-66 |
| Il1rl1 | 4.82E-65 |
| Cd81 | 9.43E-59 |
| Gata3 | 2.70E-45 |
| Ltb4r1 | 3.69E-45 |
| Cysltr1 | 3.73E-43 |
| Arg1 | 4.02E-41 |
| Slc7a8 | 1.32E-40 |
| Ccdc184 | 2.55E-40 |

Th17 Cells

| Gene | P_adj |
|---|---|
| Cd163l1 | 5.42E-83 |
| Tcrg-V6 | 1.71E-82 |
| Blk | 8.14E-79 |
| Rorc | 1.13E-64 |
| Kcnk1 | 2.72E-64 |
| Aqp3 | 5.93E-64 |
| Actn2 | 3.92E-61 |
| Sox13 | 4.85E-54 |
| Serpinb1a | 1.41E-52 |
| Ly6g5b | 2.54E-50 |

NK/NKT Cells

| Gene | P_adj |
|---|---|
| Fcer1g | 2.81E-71 |
| Klrb1c | 9.32E-55 |
| Cd7 | 6.33E-53 |
| Tyrobp | 7.32E-52 |
| Xcl1 | 2.03E-50 |
| Ncr1 | 1.61E-48 |
| Klrd1 | 1.95E-39 |
| Klrk1 | 3.62E-37 |
| Klre1 | 5.70E-36 |
| Cd160 | 5.71E-32 |

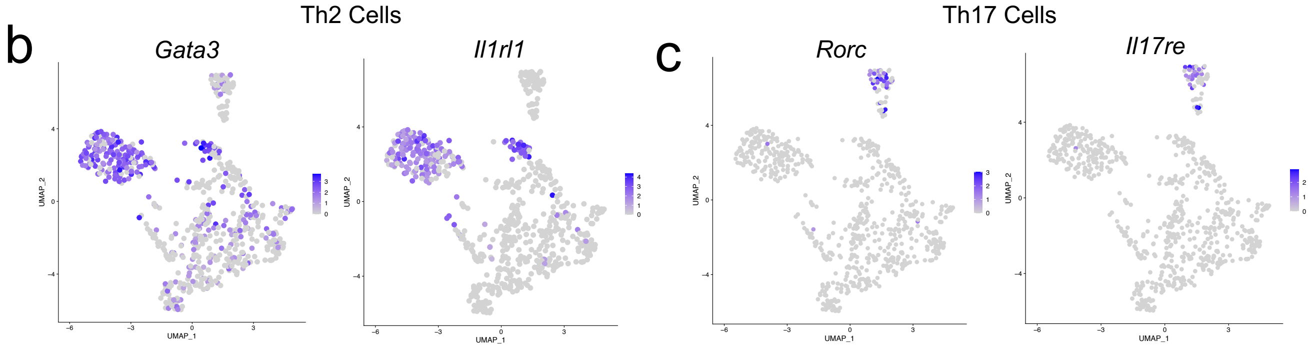


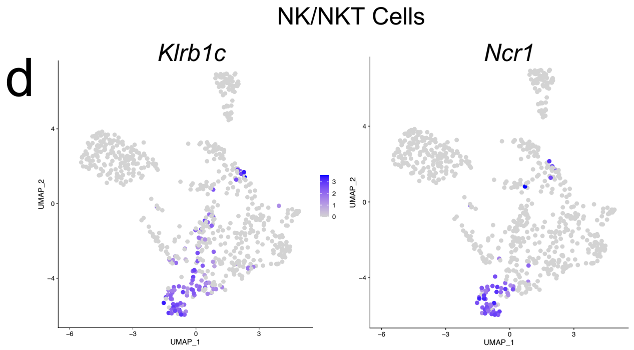


Supplementary Figure 5

a

Mature B Cells

| Gene | P_adj |
|---|---|
| Ms4a1 | 4.55E-85 |
| Ly6d | 2.14E-47 |
| Tmsb4x | 5.20E-42 |
| Cd79a | 9.76E-42 |
| Pld4 | 3.20E-38 |
| Cd79b | 4.31E-38 |
| Hck | 2.69E-37 |
| Iglc1 | 5.42E-36 |
| Crip1 | 6.35E-32 |
| Ptpn6 | 5.86E-30 |

Activated B Cells

| Gene | P_adj |
|---|---|
| H2-Aa | 6.73E-85 |
| H2-Eb1 | 4.71E-84 |
| Rps24 | 1.96E-71 |
| H2-Ab1 | 2.82E-67 |
| Rps27 | 5.73E-62 |
| Shisa5 | 6.17E-62 |
| Rps21 | 9.44E-62 |
| Sell | 1.99E-60 |
| Gimap3 | 3.40E-57 |
| Gimap4 | 4.33E-55 |

Immature B Cells

| Gene | P_adj |
|---|---|
| Rag1 | 7.91E-81 |
| Xrcc6 | 1.88E-80 |
| Smarca4 | 2.09E-71 |
| Atp1b1 | 2.64E-71 |
| Tifa | 2.30E-70 |
| Dnajc7 | 1.39E-65 |
| Arl5c | 1.69E-65 |
| Il7r | 1.14E-64 |
| Rag2 | 9.65E-53 |
| Fam53b | 3.20E-51 |

Proliferating Cells

| Gene | P_adj |
|---|---|
| Ccnd2 | 3.11E-31 |
| Srm | 4.07E-29 |
| Mettl1 | 2.00E-27 |
| Myc | 2.83E-21 |
| Apex1 | 3.94E-21 |
| Wnt10a | 4.34E-19 |
| Nop16 | 5.69E-19 |
| Grwd1 | 1.38E-18 |
| Bcl3 | 3.79E-18 |
| Mrto4 | 3.33E-17 |

Activated B Cells

b

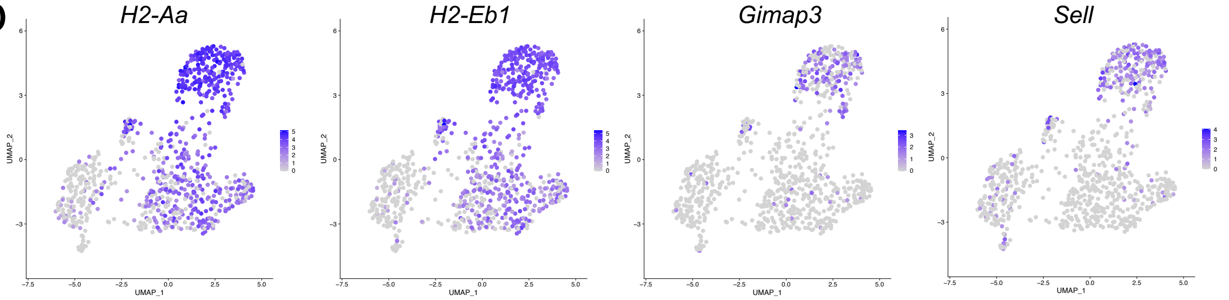


Immature B Cells

c

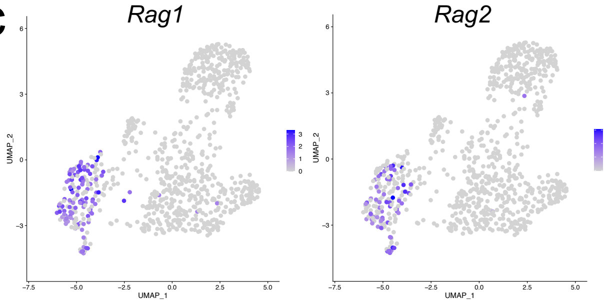


Proliferating Cells

d

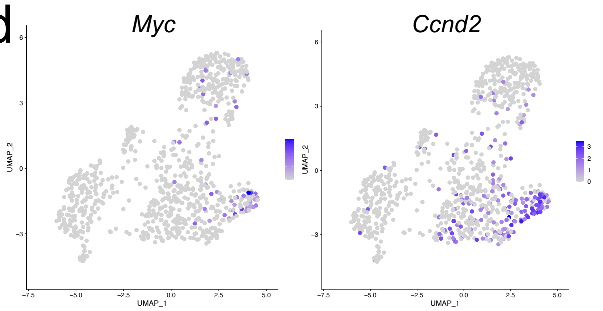


Supplementary Figure 6

| Cell Type | Cell Counts Sham | Cell Counts TBI |
|---|---|---|
| Activated Macs 1 | 185 | 499 |
| Activated Macs 2 | 199 | 319 |
| Macs 3 | 24 | 31 |
| B Cells 1 | 217 | 172 |
| B Cells 2 | 172 | 137 |
| Immature/Diff B Cells | 63 | 92 |
| CD3+ T Cells | 150 | 240 |
| Activated T Cells | 148 | 169 |
| NK Cells | 68 | 55 |
| Dendritic Cells | 131 | 200 |
| Plasmacytoid Dendritic Cells | 18 | 42 |
| Neutrophils | 16 | 26 |
| Proliferating Cells | 36 | 58 |
| Fibroblasts | 124 | 781 |
| Endothelial Cells 1 | 235 | 380 |
| Endothelial Cells 2 | 78 | 199 |
| Pericytes | 20 | 69 |
| Choroid Plexus | 58 | 135 |
| Pineal Gland Cells | 31 | 75 |
| Stem Cells | 35 | 47 |
| Clotting Related | 34 | 43 |

Table 1
